# Electrostatic-assemblies of single-walled carbon nanotubes and sequence-tunable peptoid polymers detect a lectin protein and its target sugars

**DOI:** 10.1101/500611

**Authors:** Linda Chio, Jackson Travis Del Bonis-O’Donnell, Mark A. Kline, Jae Hong Kim, Ian R. McFarlane, Ronald N. Zuckermann, Markita P. Landry

## Abstract

A primary limitation to real-time imaging of metabolites and proteins has been the selective detection of biomolecules that have no naturally-occurring or stable molecular recognition counterparts. We present developments in the design of synthetic near-infrared fluorescent nanosensors based on the fluorescence modulation of single-walled carbon nanotubes (SWNT) with select sequences of surface-adsorbed *N*-substituted glycine peptoid polymers. We assess the stability of the peptoid-SWNT nanosensor candidates under variable ionic strengths, protease exposure, and cell culture media conditions, and find that the stability of peptoid-SWNTs depends on the composition and length of the peptoid polymer. From our library, we identify a peptoid-SWNT assembly that can selectively detect lectin protein wheat germ agglutinin (WGA) with a sensitivity comparable to the concentration of serum proteins. This WGA protein nanosensor is characterized with near-infrared spectroscopy and microscopy to study protein-nanosensor interaction parameters. To demonstrate the retention of nanosensor-bound protein activity, we show that WGA on the nanosensor produces an additional fluorescent signal modulation upon exposure to the lectin’s conjugate sugars, suggesting the lectin protein selectively binds its target sugars through ternary molecular recognition interactions relayed to the nanosensor. Our results inform design considerations for developing synthetic molecular recognition elements by assembling peptoid polymers on SWNTs, and also demonstrate these assemblies can serve as optical nanosensors for lectin proteins and their target sugars. Together, these data suggest certain peptoid sequences can be assembled with SWNTs to serve as versatile optical probes to detect proteins and their molecular substrates.

---

Single-walled carbon nanotubes (SWNTs) have emerged as promising signal transduction elements for molecular imaging owing to their relatively tissue-transparent optical properties, photostability, low toxicity when functionalized, and nanometer size.^1–3^ SWNTs exist in many distinct chiralities described by the chiral vector of the carbon lattice which dictates the SWNT fluorescence excitation and emission wavelength, making SWNTs suitable for ratiometric sensing.^4,5^ SWNTs notably fluoresce in the near-infrared (NIR), a region of the electromagnetic spectrum following visible wavelength photon scattering by biological tissues, but prior to absorption of photons by water.^6,7^ This NIR window exhibits minimal photon attenuation, suitable for deep tissue bio-imaging applications.^8^ However, pristine SWNTs exhibit a hydrophobic π-conjugated surface lattice and require polymer or surfactant encapsulation for both colloidal stability and for exciton radiative recombination-based NIR fluorescence. The encapsulation of SWNTs with biomolecules and biopolymers can serve a dual purpose of conferring biocompatibility, but can also enable molecular recognition for biological analytes of interest. In particular, SWNTs can be functionalized to be selective and sensitive optical nanosensors via surface adsorption by polymers with a method known as corona phase molecular recognition (CoPhMoRe).^9^ In this technique, polymers are electrostatically pinned to the surface of the SWNT and adopt a particular conformation that can recognize a small molecule via selective modulation of the SWNT exciton recombination rate (intensity change), or band-gap (wavelength shift).^10^ To date, polymer-SWNT nanosensors have been created with DNA oligomers, peptides, and fluorescein-, rhodamine-, or carbodiimide-derived polymers as the nanosensor corona phase.^11–16^ These polymer-SWNT nanosensors have been successful in the detection of DNA hybridization, neurotransmitters, and vitamins. However, detection of proteins remains a challenging task, owing to the complexity of these larger analytes and the limited chemical stability of the molecular recognition elements often used for protein detection such as antibodies. Thus, SWNT CoPhMoRe is an attractive synthetic platform for optical detection of protein analytes, with Rap1, HIV-1, fibrinogen, and insulin protein SWNT nanosensors developed recently.^12,17,18^

Of the polymers leveraged for SWNT nanosensor design, biopolymers such as polynucleotides and peptides are often preferred owing to structural tunability with which they can be synthesized or produced by bio-organisms. However, polynucleotides and peptides are susceptible to enzymatic degradation by nucleases or proteases, respectively, and as such their conditional stability limits long-term use in complex biological systems. Conversely, synthetic polymers may require lengthy synthesis and are difficult to control in terms of sequence, length dispersity, and structural tunability. Furthermore, both biopolymers and synthetic polymers are limited by a lack of monomer sets for creating chemical diversity. Therefore, future advancements in the area of synthetic protein nanosensors will require generation of biomimetic polymers amenable to facile synthesis, with a large monomer space, which are also resistant to enzymatic degradation. To this end, herein we synthesize a library of peptoids, *N*-substituted glycine polymers, to serve as protein molecular recognition elements when adsorbed on SWNT surfaces. Peptoids are sequence-defined synthetic polymers that draw inspiration from peptides, the building blocks of proteins, with synthesis that is amenable to robotic automation and a large monomer space of primary amines.^19^ Peptoids are created through stepwise solid phase submonomer synthesis with high sequence specificity and can include a wide variety of non-proteinogenic chemical functionalities such as alkynes, glycosylation, and fluorophores.^20–22^ Peptoids are resistant to proteases and can remain stable in the body for day-long timescales.^23^ Recently, peptoids have been shown to self-assemble into supramolecular nanosheets that are capable of specific multivalent interactions with enzymes and proteins such as kinases, lectins, and Shiga toxin.^24,25^ Because peptoids are designable and tunable proteomimetic materials, they are well suited to address the current need of chemical diversity of SWNT polymer coronas.^26^

We present the development of peptoid polymer-SWNT (peptoid-SWNT) assemblies and their characterization for implementation as protein nanosensors with UV-Vis-NIR absorbance, atomic force microscopy, NIR spectroscopy and imaging. We first investigate the primary interactions of the peptoid polymer with SWNT, and present our findings on the stability of these peptoid-SWNT assemblies upon exposure to long-term laser illumination, variable solution ionic strength conditions, complex biocompatible media, and proteases. We next show that certain peptoid-SWNT assemblies can serve as nanosensors through secondary interactions to enable selective and sensitive WGA lectin protein detection. We further show that these peptoid-SWNT nanosensors have the capacity for ternary analyte interaction and detection, where the WGA protein tethered to the SWNT nanosensor can in turn detect WGA’s target sugars. Our work informs key design parameters for the noncovalent adsorption of peptoids with SWNT and their development as molecular recognition elements for protein sensing and ternary analyte interactions.

## Peptoid Polymer Design for Surface Adsorption to SWNT

Peptoid polymers are created with an automated submonomer approach that provides ease of synthesis and control over polymer length and sequence (Scheme 1).^27^ The chemical sequence of the peptoid is dictated by the selective order in which variable side chain groups are added to the growing peptoid chain via primary amines.

**Scheme 1:**
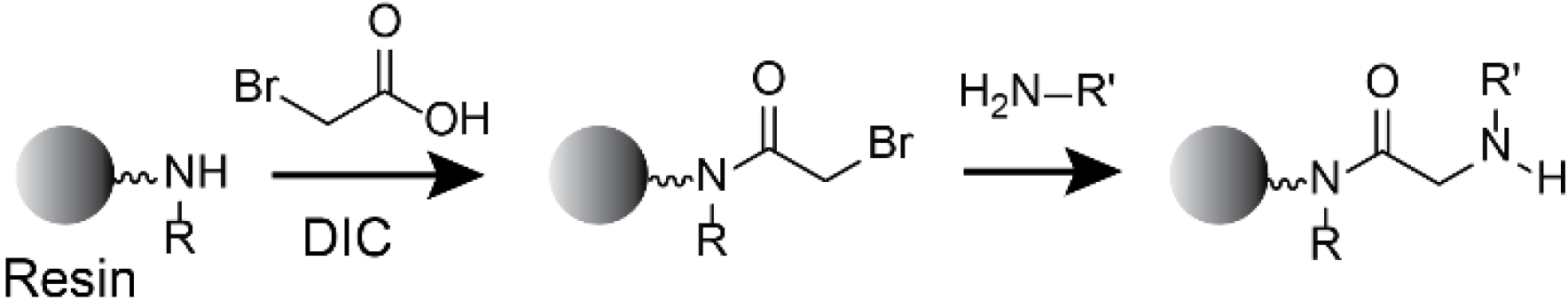
Peptoid synthesis with the solid-phase two-step submonomer method. Peptoids are synthesized through acylation using bromoacetic acid and, *N,N*’-diisopropylcarbodiimide (DIC) followed by displacement with a primary amine.

To gauge the possibility of creating peptoid-SWNT assemblies for use as protein nanosensors, we first elucidated the peptoid design parameters necessary for rendering colloidally stable peptoid-SWNT assemblies. We synthesized a small library of 11 amphiphilic peptoid structures, outlined in Table 1 (see SI. 1 for full chemical structures), of variable sequences, charges, polarities, and lengths to probe peptoid-SWNT assembly efficacy using nonpolar monomers: *N*-(2-phenylethyl)glycine (Npe), *N*-isoamylglycine (Nia), and *N*-phenylglycine (Nph); uncharged hydrophilic monomers: *N*-(2-methoxyethyl)glycine (Nme) and *N*-2-(2-(2-methoxyethoxy)ethoxy)ethylglycine (Nte); positively charged *N*-(2-aminoethyl)glycine (Nae); and negatively charged *N*-(2-carboxyethyl)glycine (Nce). We included several diblock peptoids similar to previous antibody-mimetic nanosheet forming peptoids such as Block36 (Nae-Npe)9-(Nce-Npe)9, DB1 (Nae-Nia)7-(Nce-Nia)7, and DB2 (Nae-Nph)4-(Nce-Nph)4.^28^ We synthesized several polar but uncharged peptoid polymers such as Pep1 (Nte-Npe)14, Pep3 (Nme-Npe)14, as well as a hybrid charged and uncharged polar polymer Pep2 (Nte-Npe-Nce-Npe)7. We included a positively charged polymer, PA28 (Nae-Npe)14, and negatively charged polymers of alternating Npe and Nce (PC) monomers with final polymer lengths of 18, 28, and 36 (PC18, PC28, and PC36, respectively), an anchor-loop peptoid consisting of a (Nce-Npe)9 ‘anchor’ peptoid sequence that can adsorb to the carbon nanotube flanking a 6-monomer loop consisting of 3 *N*-butylglycine (Nbu) and 3 *N*-(*N*’-pyrrolidinonylpropyl)glycine (Npp) monomers ((Nce-Npe)9-Nbu-Nbu-Npp-Npp-Nbu-Npp-(Nce-Npe)9, and further abbreviated as ProLoop). The loop segment within the ProLoop peptoid interacts semi-selectively with wheat germ agglutinin (WGA), a 36 kDa lectin, a sugar-binding protein, as assessed with a FRET binding assay (see supplemental information figure 2 for selectivity and sensitivity FRET screen results).

**Table 1:**
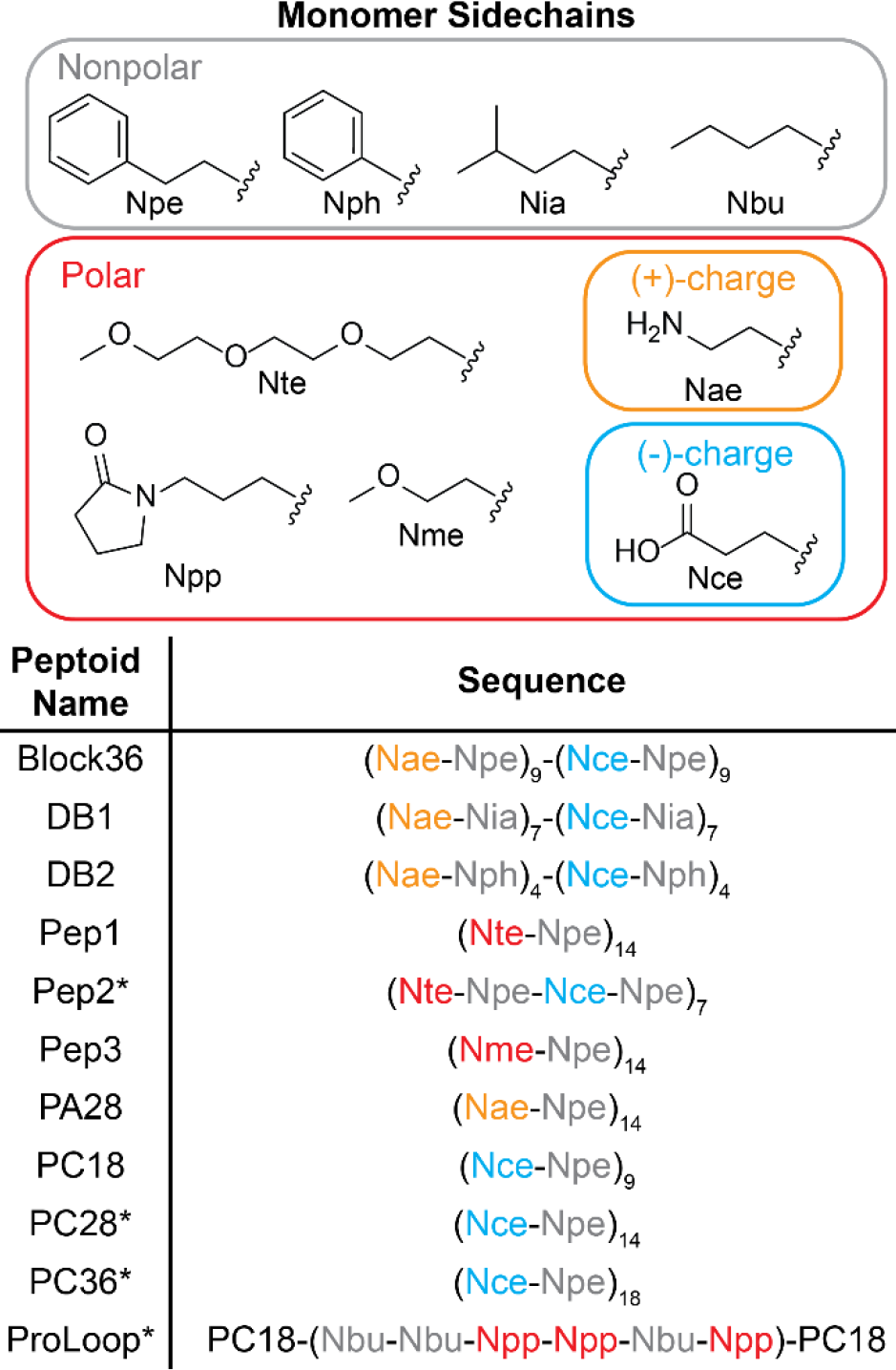
Library of 11 peptoid structures tested for adsorption to SWNT. Peptoids denoted with an asterisk (*) successfully adsorb on SWNT to form peptoid-SWNT assemblies; while other peptoids are unable to suspend SWNT.

| Monomer Sidechains |  |
| --- | --- |
| Nonpolar |  |
| Polar |  |
| Peptoid Name | Sequence |
| Block36 | (Nae-Npe)<sub>9</sub>-(Nce-Npe)<sub>9</sub> |
| DB1 | (Nae-Nia)<sub>7</sub>-(Nce-Nia)<sub>7</sub> |
| DB2 | (Nae-Nph)<sub>4</sub>-(Nce-Nph)<sub>4</sub> |
| Pep1 | (Nte-Npe)<sub>14</sub> |
| Pep2* | (Nte-Npe-Nce-Npe)<sub>7</sub> |
| Pep3 | (Nme-Npe)<sub>14</sub> |
| PA28 | (Nae-Npe)<sub>14</sub> |
| PC18 | (Nce-Npe)<sub>9</sub> |
| PC28* | (Nce-Npe)<sub>14</sub> |
| PC36* | (Nce-Npe)<sub>18</sub> |
| ProLoop* | PC18-(Nbu-Nbu-Npp-Npp-Nbu-Npp)-PC18 |

The peptoid library was evaluated for each peptoid’s ability to adsorb to SWNTs. Peptoid-SWNT adsorption was accomplished with probe-tip sonication (see Methods) and adsorption efficacy was confirmed with UV-Vis-NIR absorbance spectroscopy and NIR fluorescence spectroscopy (Fig. 1, SI. 3). UV-Vis-NIR absorbance was used to measure the optical density, from which yield can be calculated (see Methods), of colloidally stable peptoid-SWNT assemblies. NIR fluorescence spectroscopy further confirms peptoid adsorption as only stable peptoid-SWNT assemblies exhibit NIR fluorescence. Probe-tip sonication was performed in 50 mM borate buffer (pH 9.2) and peptoid-SWNT assemblies were diluted in phosphate buffered saline (PBS, pH 7.4) prior to fluorescence measurements, to represent physiological conditions in preparation for downstream biological applications in protein sensing.

**Figure 1:**
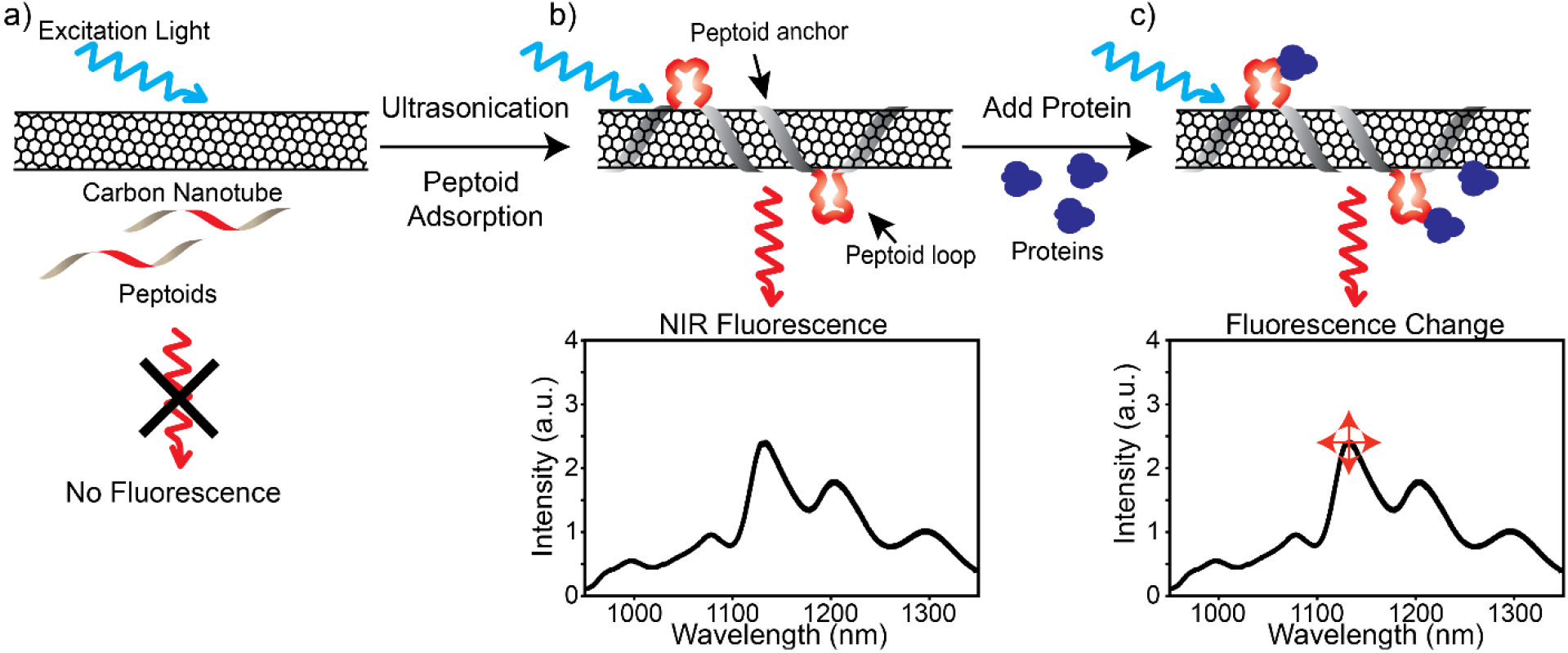
Platform for peptoid-SWNT assembly. a) No NIR fluorescence is observed prior to adsorption of peptoids to SWNT surfaces. b) Ultrasonication promotes the formation of peptoid-SWNT assemblies with a distinct NIR fluorescence spectrum that is c) modulated by the addition of a target protein (blue) that binds to the protein-recognition loop (red).

We found that four of the eleven peptoids stably adsorb onto SWNTs, with successful peptoid-SWNT assemblies labeled with an asterisk (*) in Table 1. Adsorption efficiency was compared by measuring the UV-Vis-IR absorbance spectrum of the assembly, where a higher absorbance is indicative of higher peptoid-SWNT yield (SI 3). We found that peptoid polymer length affects the effectiveness of peptoid-SWNT adsorption. To assess the effect of peptoid length on peptoid-SWNT stability, we compared peptoids comprised of the same carboxyethyl-phenethyl repeat with lengths of 18, 28, and 36 monomers. We found the peptoid-SWNT adsorption efficiency as determined from peptoid-SWNT yields is roughly proportional to peptoid length: PC36 (yield = 165.1 mg/L) ≈ PC28 (194.9 mg/L) > PC18 (9.2 mg/L). We posit that longer peptoid polymers adsorb more strongly to SWNTs owing to the increased number of contacts made between a longer peptoid polymer and the surface of the SWNT. We also found that peptoid hydrophilicity is an important contributor to peptoid-SWNT adsorption efficiency. Pep1, Pep2, Pep3, and PC28 peptoids are of the same 28 monomer length, with alternating monomers of aromatic phenethyl and hydrophilic monomers of either triethyloxy, monoethoxy, or carboxyethyl. We show that the efficiency of peptoid adsorption to SWNT follows the trend of PC28 > Pep2 > Pep1 ≈ Pep3, with Pep1 and Pep3 unable to suspend SWNT, and emerges from the trend of peptoid hydrophilicity as observed by HPLC (SI. 4). Lastly, we note that peptoid charge also facilitates peptoid-SWNT colloidal stabilization: both the peptoid backbone and the SWNT are nonpolar and hydrophobic, and require hydrophilicity by charge or polarity to create stable peptoid-SWNT assemblies in aqueous buffer. Of the charged 28-mer peptoids, the highly negatively charged PC28 best adsorbed to SWNT with a yield of 194.9 mg/L, whereas the less charged Pep2 adsorbed least with a yield of 19.8 mg/L and the neutrally charged Pep1 failed to adsorb onto SWNT. Therefore, we find that charged monomers can confer colloidal stabilization of SWNT.

Several peptoids did not adsorb to SWNT. Peptoids lacking aromatic hydrophobic residues, such as DB1, did not suspend SWNT presumably due to a lack of π-π interactions. Conversely, peptoids with aromatic groups such as Block36 have previously been shown to self-assemble into supramolecular nanosheets driven by zwitterionic stabilizing interactions of the charged groups, hydrophobic interactions, and π-π stabilization between the aromatic rings of the peptoid.^28^ Thus, when Block36 is adsorbed to SWNTs, we propose that peptoid zwitterionic and hydrophobically-driven self-interactions dominate over peptoid-SWNT interactions, resulting in largely unstable SWNT assemblies in favor of spontaneous formation of peptoid nanosheets as confirmed by AFM (SI. 5). DB2, not known to form nanosheets, also did not adsorb to SWNT, presumably due to the larger steric hinderance of phenyl monomers compared to the 2-phenylethyl monomers.

From our initial screen, we identified PC36 as a highly stable peptoid for SWNT stabilization, and utilized this ‘anchor’ for peptoid adsorption to SWNT (SI. 3). We also identified ProLoop as a stable peptoid for SWNT adsorption with a moderate yield of 103.1 mg/L. Additionally, the 6-monomer central segment of the ProLoop peptoid has demonstrated binding affinity for lectin protein WGA as measured by FRET (SI. 2). Thus, the ProLoop-SWNT assembly, and the PC36-SWNT assembly, were selected for downstream stability characterization and for use as fluorescence WGA protein nanosensors.

## Peptoid-SWNT Nanosensor Fluorescence Stability

To validate peptoid-SWNT assemblies as fluorescent protein nanosensors, we next sought to demonstrate the stability of peptoid-SWNT assemblies in a range of conditions expected for bioimaging applications. We first examined the effect of salt and buffer conditions, as such parameters have been shown to affect the stability of SWNT assemblies.^29–31^ We also examined the stability of the SWNT assemblies to continuous laser exposure and protease activity.

To test salt stability, peptoid-SWNT assemblies were incubated overnight in sodium chloride (NaCl) solutions ranging in concentration from 1 mM to 500 mM, and peptoid-SWNT fluorescence was subsequently measured (Fig. 2a). PC28 and PC36 peptoid-SWNT assemblies remained colloidally stable and exhibited NIR fluorescence under a 1 mM to 500 mM NaCl range of ionic strengths. NaCl concentrations higher than 500 mM were shown to destabilize the peptoid-SWNT assemblies and induce SWNT aggregation. Pep2-SWNT assemblies were unstable in NaCl solutions with concentrations higher than 10 mM and formed visible aggregates at higher ionic strength (SI. 6). Recent work demonin lower SWNT fluorescence (Fig. 2b).^30^ Additionally, we tested the effect of salt composition on the fluorescence of peptoid-SWNT assemblies. Recently, divalent salts have been shown to induce a wavelength shift in the fluorescence spectra of DNA-SWNT assemblies. In these studies, it is hypothesized that these solvatochromic shifts are due to induced conformational changes in the backbone a DNA polymer along the SWNT, and correlated with the stiffness of the polymer backbone.^31^ We observed significant wavelength shifts in the fluorescence spectra of PC28 and PC36 peptoid-SWNT assemblies upon addition of CaCl2 salt, and minor shifts in the fluorescence spectra of ProLoop and Pep2 peptoid-SWNT assemblies (SI. 7), suggesting that multiple factors, of which salts may be but one, affect peptoid flexibility and thus binding stability on SWNT.

**Figure 2:**
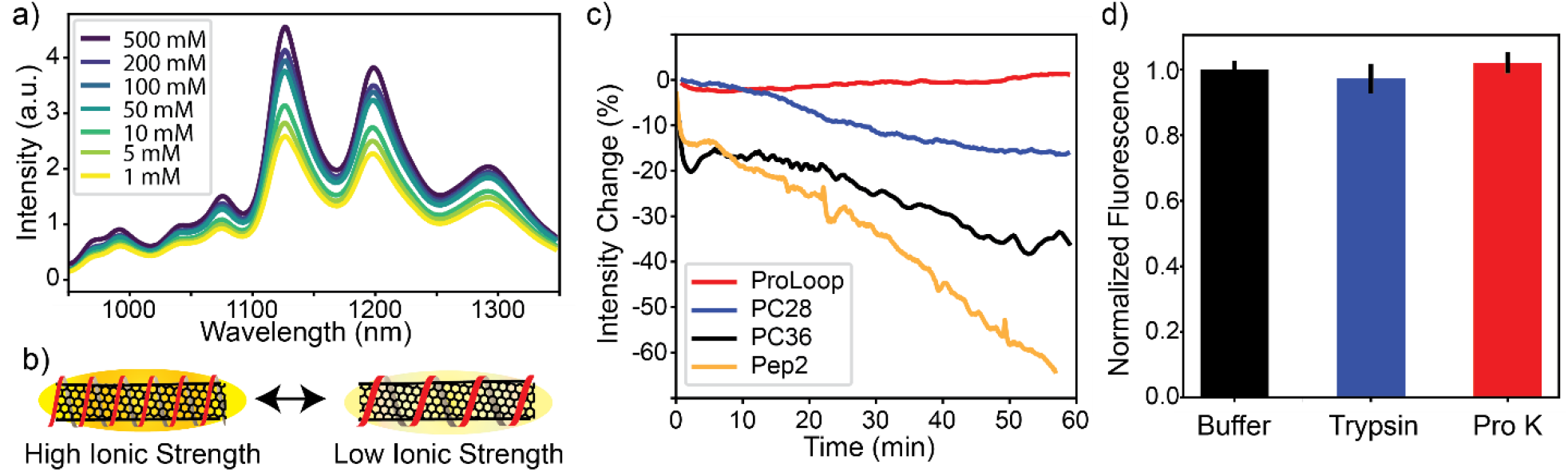
Peptoid-SWNT stability tests. a) Peptoid-SWNT fluorescence, shown here for PC36-SWNT, increases with increasing ionic strength, suggesting b) peptoid surface coverage of SWNT increases with ionic strength. c) Fluorescence of peptoid-SWNT assemblies under continuous laser irradiation shows ProLoop-SWNT as the most stable assembly in this study. d) ProLoop-SWNT exposure to trypsin or proteinase K (Pro K) for 24 hours reveals no loss of fluorescence. Error bars denote standard deviation.

We next subjected each peptoid-SWNT assembly to an hour-long imaging experiment to probe peptoid-SWNT fluorescence intensity and thus stability to continuous laser exposure. Excitation of peptoid-SWNT samples by a 77 mW 721 nm laser for SWNT spectroscopy stabilizes to a temperature of 37°C in the sample well within 1 minute of laser irradiation (SI. 8). Peptoid-SWNT assemblies did not show wavelength shifts during the hour-long spectral measurement though some showed chirality-independent changes in fluorescence (SI. 9). Our results showed that the ProLoop-SWNT construct exhibits the most stable fluorescence with a negligible 2.1 ± 6.6 % (mean ± standard deviation (SD)) decrease in fluorescence after 1 hour of laser illumination (Fig. 2c). In contrast, PC28 and PC36 both showed moderate fluorescence perturbations of 12.2 ± 8.9 % (mean ± SD) and 36.8 ± 17.8 % (mean ± SD) decrease in fluorescence, respectively, while Pep2-SWNT showed the largest fluorescence perturbation with a 69.7 ± 27.1% (mean ± SD) decrease in fluorescence. We note that peptoid-SWNT stability trends observed herein follow trends in peptoid-SWNT adsorption efficiency, although the ProLoop-SWNT exhibited moderate adsorption efficiency and highest stability. The superior fluorescence stability and the incorporation of a WGA molecular recognition loop sequence led us to choose the ProLoop-SWNT assembly as an exploratory candidate for use as an optical nanosensor to detect the presence of WGA protein. Additionally, we tested the stability of the ProLoop-SWNT nanosensor against two proteases: trypsin and proteinase K. These proteases were chosen for their ability to digest a wide range of peptide sequences, specifically positively charged, nonpolar, and aliphatic amino acids. We found that 24 hour incubation of peptoid-SWNT in proteases yielded minimal loss of fluorescence and hence no significant peptoid proteolysis (Fig. 2d, SI. 10).^32^

## ProLoop-SWNT Nanosensors Exhibit Fluorescence Modulation in Response to Wheat Germ Agglutinin Protein

The ProLoop-SWNT assembly was assessed for use as a fluorescent nanosensor for WGA protein, a sugar-binding lectin, via binding of WGA to the ProLoop peptoid sequence (SI. 2). ^33^ ProLoop-SWNT nanosensors were diluted to a concentration of 5 mg/L and the baseline fluorescence was measured (denoted as *Io*). WGA was added to the nanosensor and subsequent fluorescence (denoted as *I*) was measured at consecutive timepoints. We found that the ProLoop-SWNT nanosensor exhibited a decrease in fluorescence upon the introduction of WGA. The fluorescent response was time-dependent and reached equilibrium within 60 minutes following addition of 10 μM WGA (Fig. 3a). We next characterized the ProLoop-SWNT nanosensor sensitivity to WGA through a concentration screen of 0.1 μM to 14.9 μM WGA. We measured the ProLoop-SWNT nanosensor change in fluorescence 60 minutes following protein addition. Assuming a single protein binding to each peptoid binding site, we modeled the equilibrium binding of ProLoop-SWNT with an enzyme-substrate binding model, as denoted by the red curve on the concentration plot (Fig. 3b, SI. 13).

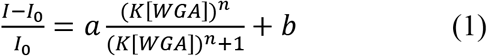

In equation 1, *I* denotes the ProLoop-SWNT nanosensor intensity after 60 minutes, *Io* denotes the initial nanosensor intensity at 0 minutes, [*WGA*] the concentration of WGA in the system, K the equilibrium binding constant, and *a* and *b* denote scaling factors (full model derived in SI. 13). Since WGA is composed of two symmetric monomeric units, we assumed the Hill coefficient (n), which denotes the cooperativity of the analyte bound, to be 2 for the fit to converge (goodness of fit of the model is discussed in SI. 13).^34^ Using nonlinear least-squares fitting, the model parameters were found to be a = −15.72, K = 0.0331 μM^-1^, and b = 0.068. With this equilibrium model of WGA binding to ProLoop-SWNT, we calculated the limit of detection of ProLoop-SWNT to WGA as 3.4 μM (SI. 13), a concentration comparable to the average serum concentration of common blood proteins such as albumin (600 μM), IgG (100 μM), and fibrinogen (7.5 μM).^35^ Interactions between the ProLoop-SWNT nanosensor and WGA protein was further confirmed by atomic force microscopy showing WGA proteins attached to the surface of the nanosensor (SI.12).

**Figure 3:**
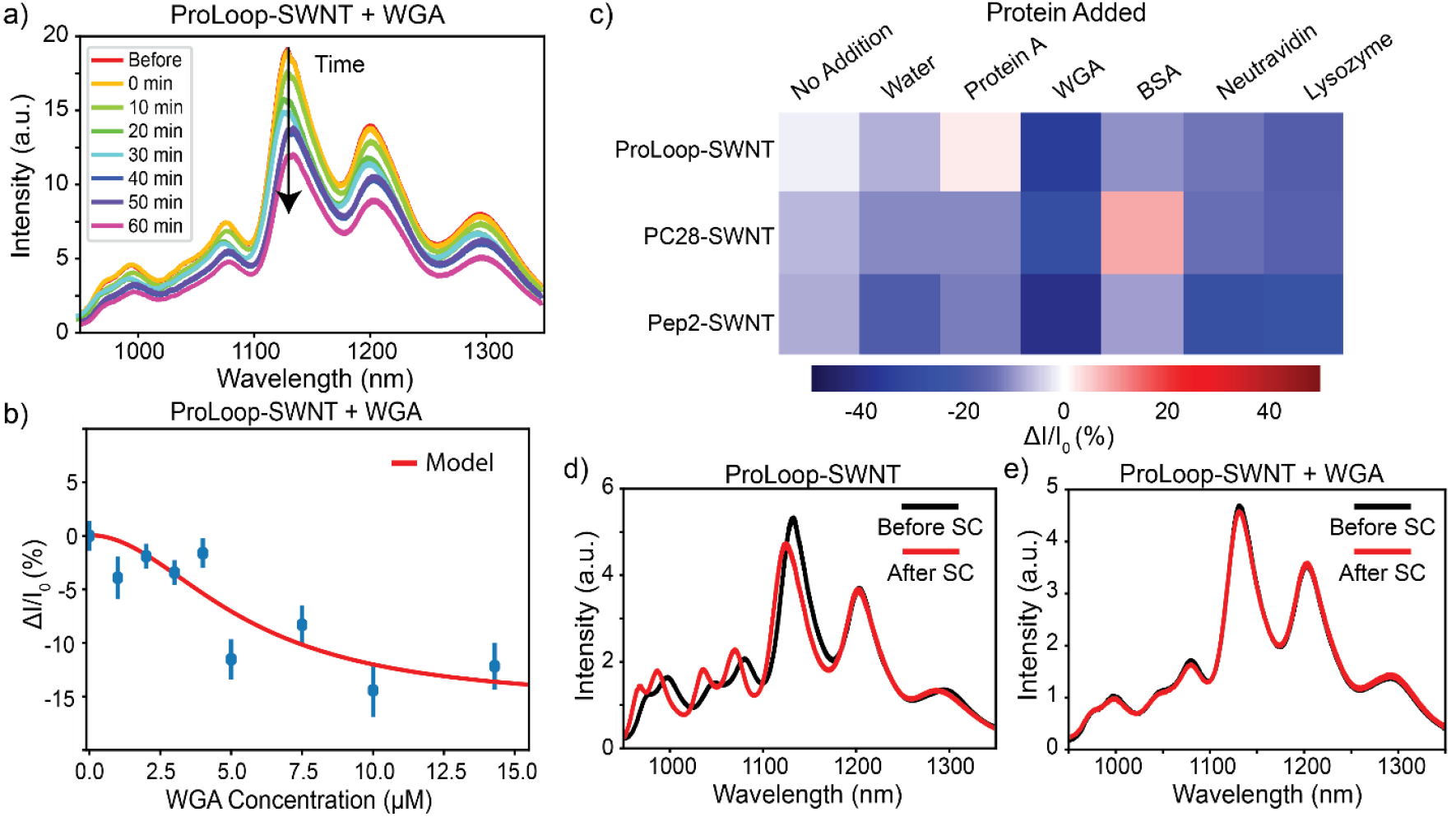
ProLoop-SWNT nanosensor characterization. a) ProLoop-SWNT exhibited a decrease in fluorescence upon addition of WGA protein. b) The decrease in fluorescence of ProLoop-SWNT was most selective for WGA against other proteins. c) The equilibrium decreases in fluorescence of ProLoop-SWNT as a function of WGA concentration yields a 3.4 μM limit of WGA detection. Fluorescence change was normalized to 0 at 0 μM WGA to account for nanosensor dilution, and error bars denote standard deviation. d) Addition of sodium cholate (SC) to ProLoop-SWNT perturbed its fluorescence emission with a solvatochromic shift, indicating the surfactant can bind to SWNT by partially displacing the peptoid polymer. e) This surfactant-induced solvatochromic shift is eliminated after ProLoop-SWNT binds WGA.

Next, the selectivity of the ProLoop-SWNT nanosensor was tested by measuring the fluorescence response of ProLoop-SWNT upon exposure to bovine serum albumin (BSA), NeutrAvidin, protein A, and lysozyme proteins (Fig. 3c). Both BSA and NeutrAvidin are proteins known to bind the surface of SWNT, whereas protein A is a negatively charged protein at physiological pH, and lysozyme is a positively charged protein at physiological pH.^12,36^ This modest protein library afforded a screen for ProLoop-SWNT selectivity towards WGA. We found that ProLoop-SWNT nanosensors show the highest magnitude in fluorescence response of −34.5 ± 15.6% (mean ± SD) upon addition of 10 μM WGA, with minor fluorescence modulation upon addition of 5 mg/mL protein A or 5 mg/mL lysozyme (2.03 ± 8.2% and −17.26 ± 6.4% (mean ± SD), respectively). Conversely, PC28-SWNT and Pep2-SWNT assemblies also showed a high magnitude in fluorescence response upon addition of 10 μM WGA (−27.9 ± 18.1% and −39.5 ± 3.7% (mean ± SD), respectively). However, addition of proteins, buffer, or nothing to PC28-SWNT and Pep2-SWNT induced monotonic decreases in fluorescence, indicating that fluorescence modulation of PC28 and Pep2 constructs derives primarily from their low colloidal stability. Therefore, ProLoop-SWNT exhibited the highest selectivity towards WGA protein and superior colloidal stability over the 1-hour molecular recognition experimental window.

## Probing the Interaction Between Peptoid-SWNTs and WGA

To further understand the interactions of WGA with peptoid-SWNT assemblies and also the identified ProLoop-SWNT nanosensor, we studied the fluorescence change of the nanosensors upon surface perturbation by surfactant. Previous studies have shown that surfactants including sodium dodecyl sulfate, sodium cholate (SC), and sodium dodecylbenzenesulfonate, can bind to the exposed SWNT surface of polymer-SWNTs and exclude water from SWNTs.^15^ This water exclusion causes a change in the dielectric environment of the nanosensors and induces a solvatochromic shift that can be implemented to study the accessibility of the SWNT surface in a polymer-SWNT construct.

Addition of 0.5% (w/v) SC to ProLoop-SWNT nanosensors induced a solvatochromic blue shift in fluorescence for most nanotube chiralities, with the peaks at shorter fluorescence emission wavelengths showing the largest peak wavelength perturbation (Fig. 3d) with peak wavelength shifts ranging from 0.97 nm to 11.76 nm. This wavelength shift was also exhibited by the other peptoid-SWNT nanosensors (SI. 14). Notably, Pep2-SWNT was only minimally perturbed by SC addition, showing on average less than 1 nm shift after the addition of SC (SI. 14). This is likely due to the triethylether sidechains in Pep2 that resemble polyethylene glycol (PEG) and may limit accessibility of SC to the SWNT surface. PEG is often used as an antifouling and biocompatibility coating on biological device and nanoparticle surfaces, and antifouling peptoids have previously been synthesized to recreate this property of PEG or other antifouling polymers.^37^ Perhaps similarly, the triethylether monomers of the peptoid polymer may create an antifouling polymer brush along the surface of the Pep2-SWNT assembly. Thus, in future implementations of peptoid-SWNT nanosensors, antifouling properties may be engineered into the construct with the addition of PEG-like monomers to the peptoid.

Interestingly, addition of WGA to the ProLoop-SWNT assemblies prior to the addition of 0.5% SC does not show a solvatochromic shift, suggesting WGA stabilizes the ProLoop-SWNT assemblies against surfactant perturbation (Fig. 3e). Prior work has confirmed that this stabilization effect represents a strong and selective binding interaction between the molecular analyte and the polymer-SWNT assembly.^38^ To confirm we may attribute the additional stability provided to ProLoop-SWNT assemblies by selective binding of WGA, we showed that WGA by itself does not associate strongly to SWNT, and showed that attempted assembly of WGA with SWNT produced a highly unstable complex (SI. 15). These results suggest that the peptoid-SWNT assembly is necessary to stably bind WGA, and this binding of the target analyte further promotes surface stability of the nanosensor-protein complex.

## Peptoid-SWNT Stability in Biological Media

For biological applications, peptoid-SWNT nanosensors need to retain their ability to detect target proteins in complex biological environments. We showed that ProLoop-SWNT nanosensors retain their ability to respond to WGA in Dulbecco Modified Eagle’s Medium (DMEM), a common medium used for mammalian cell culture. Pre-incubation of the nanosensors with DMEM for an hour retained ProLoop-SWNT nanosensor responsivity to 10 μM WGA (Fig. 4a). DMEM preincubated ProLoop-SWNT nanosensors show a fluorescence decrease of 58.6 ± 8.6 % (mean ± SD) upon the addition of WGA against a baseline decrease of 29.3 ± 0.1 % (mean ± SD). Baseline decreases are caused by dilution effects and some nanosensor instability in DMEM.

**Figure 4:**
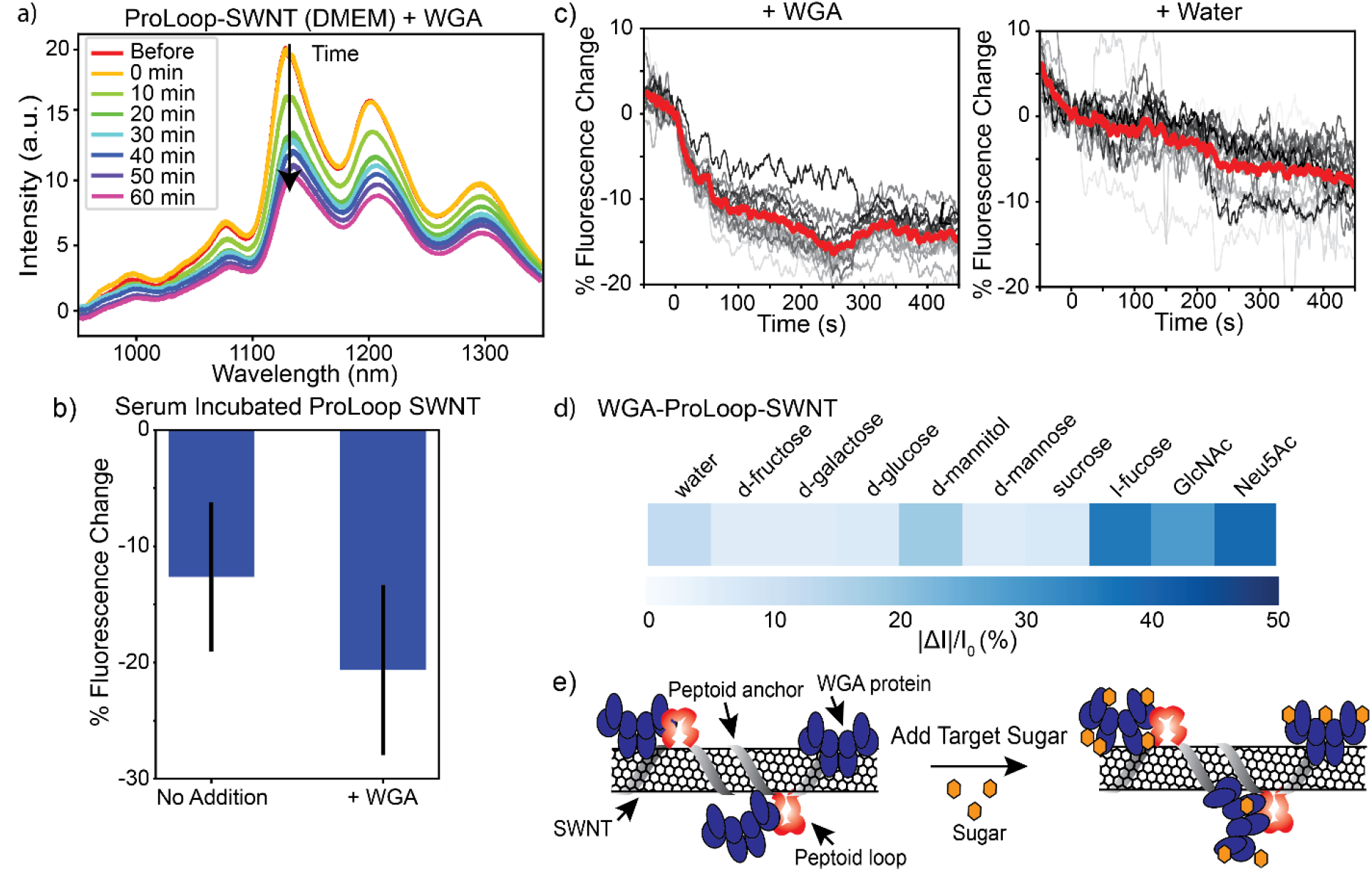
ProLoop-SWNT nanosensors for use in biological applications. a) ProLoop-SWNT is sensitive to WGA when the nanosensor is preincubated in DMEM. b) ProLoop-SWNT nanosensors show unstable fluorescence in serum, but still respond to WGA (n = 9, p < 0.05). c) Single molecule microscopy traces show that single ProLoop-SWNT nanosensors respond to WGA with a fluorescence change that is greater than baseline drift. WGA was added at t = 0 s. The gray lines plot trajectories for single nanosensors and the red line plots the average of single nanosensor intensity changes. d) WGA anchored to ProLoop-SWNT is selective for its target sugars GlcNAc and Neu5Ac, whereby ProLoop-SWNT nanosensors require the addition of both WGA and sugars to show this fluorescence signal modulation with e) a schematic illustrating possible protein rearrangements after addition of target sugars.

Lastly, we tested the responsivity of peptoid-SWNT nanosensors in serum conditions. Nanosensor response was attenuated in the presence of fetal bovine serum, with peptoid-SWNT nanosensors pre-incubated in serum for 24 hours exhibiting unstable fluorescence upon laser illumination. The fluorescence of ProLoop-SWNT pre-incubated in serum decreased by 12.6 ± 6.4% (mean ± SD) over the course of 1 hour of spectrometry without the addition of WGA, and decreased by 20.6 ± 7.3% (mean ± SD) with the addition of WGA showing the nanosensors’ ability to recognize WGA (Fig. 4b). Additional assays of ProLoop-SWNT pre-incubated in NeutrAvidin, a protein that binds bare nanotube surfaces, and recombinant human serum albumin, the most abundant protein in serum, also showed an attenuated but statistically significant fluorescence decrease response to WGA (SI. 16). These results suggest that the ProLoop-SWNT nanosensors have the capability to monitor protein dynamics in *in vitro* cell cultures, but that further optimization will be necessary to increase ProLoop-SWNT nanosensor activity in the presence of serum.

## Single-Molecule Imaging of Peptoid-SWNT Nanosensors

The fluorescence response of ProLoop-SWNT nanosensor to WGA can also be imaged with single ProLoop-SWNT nanosensors immobilized on a microscope slide, to confirm the fluorescence decrease of nanosensors are not the result of aggregation. Negatively charged ProLoop-SWNT nanosensors were surface-immobilized on a positively charged 3-aminopropyltriethoxysilane (APTES)-coated glass coverslip and imaged with NIR single nanosensor fluorescence microscopy (see Methods). Addition of WGA to a final protein concentration of 50 μM induces a fluorescence change in the saturated sensing regime of the ProLoop-SWNT nanosensor. Although single nanosensors exhibit variable fluorescence intensity trajectories over time, the average of the regions of interest show a significant decrease in fluorescence upon WGA addition −14.5 ± 2.0% (mean ± SD) compared to a control sample using only water −8.1 ± 3.5% (mean ± SD) (Fig. 4c, SI. 17). Single nanosensor imaging suggests that the molecular recognition of individual ProLoop-SWNT nanosensors can be monitored optically for dynamic biological imaging.

## WGA on Peptoid-SWNT Enables Detection of WGA’s Target Sugars

We further showed ternary nanosensor interactions whereby peptoid-SWNT nanosensors, post-interaction with WGA, can detect a secondary analyte through lectin-sugar interactions. WGA protein is a lectin, a sugar-binding protein, with specificity to two target sugars: *N*-acetylglutaminic acid (GlcNAc) and *N*-acetylneuraminic acid (Neu5Ac).^39^ We tested the activity of the protein when it is bound to the surface of ProLoop-SWNT. We first incubated ProLoop-SWNT in 10 μM WGA for an hour, before measuring its fluorescence spectrum. A panel of sugars commonly bound by lectins were tested including fructose, galactose, glucose, mannose, and fucose and complex sugars such as sucrose and mannitol in addition to the target sugars GlcNAc and Neu5Ac. The addition of the target sugars to WGA-saturated ProLoop-SWNT yielded a significant change in nanosensor fluorescence intensity (Fig. 4d). This modulation of SWNT fluorescence intensity was not observed with other sugars that other lectins are known to bind including fructose, galactose, and glucose. Interestingly, we did observe an off-target response of the WGA-saturated ProLoop-SWNT to fucose sugar. In control experiments, we confirmed that WGA and ProLoop are both required and responsible for sugar-induced SWNT fluorescence modulation (SI. 18). We posit that sugar binding of the protein causes a protein-conformation change or a perturbation to the corona phase of the ProLoop-SWNT (Fig. 4e). Furthermore, we attributed this selective fluorescence response to the 6-monomer loop in the ProLoop peptoid, since WGA-incubated with PC36-SWNT, which lacks this loop segment, does not respond to GlcNAc or Neu5Ac (SI. 19). A common challenge in protein adsorption to nanoparticles for biosensing, enzymology, and protein delivery is a reduction in protein structure and activity.^40,41^ Notably, the specificity of WGA towards its target sugars suggests WGA remains active while tethered to the SWNT surface through ProLoop-SWNT. These results further suggest peptoid-SWNT assemblies can enable detection of both proteins and also their target molecular binders.

In summary, we present design principles and validate the modular platform for the assembly of protein-like *N*-substituted synthetic glycine peptoid polymers with SWNTs. We demonstrated that certain peptoid sequences, namely ProLoop, exhibit the necessary stability to assemble on SWNT and exhibit robust photostability under conditions necessary for real-time imaging of proteins including variable buffer conditions, continuous laser illumination, and resistance to protease degradation. We further showed that ProLoop-SWNT can optically detect lectin protein WGA, and can also selectively detect the lectin conjugate sugars.

An outstanding question in nanosensor development is whether molecular recognition elements can be rationally designed for analytes of interest with synthetically-tractable polymers. To this end, including peptoids in the repertoire of polymers for suspending SWNT realizes a new space of biomimetic polymers for the development of SWNT-based nanosensors. Peptoid polymers are resistant to protease degradation, leverage facile synthesis through automated solid phase synthesis, and exhibit a large monomer space that is unavailable for conventional biopolymers. Our results suggest peptoid polymers are promising candidates for modular design of synthetic molecular recognition elements with nanomaterial substrates to serve as signal transduction elements.^19,21,42,43^

Lastly, our work shows that peptoid-SWNT nanosensors can interact with proteins while preserving the protein’s inherent activity. We demonstrated that ProLoop-SWNT can bind WGA and can fluorescently respond to the two WGA target sugars GlcNAc and Neu5Ac, demonstrating that the WGA protein remains active towards its target sugars even when bound to a peptoid-SWNT nanosensor. Despite prior reports of decreased or abolished protein activity upon protein adsorption to nanoparticles such as SWNT, it is possible that the maintenance of protein activity is due to the ProLoop-mediated binding of WGA, in lieu of WGA binding to the SWNT directly. Our results suggest peptoid polymers may serve a dual purpose of tethering proteins to SWNT in a manner that enables protein detection and also preservation of the protein’s endogenous activity for future applications in surface science, enzymology, and protein delivery.

## Supporting Information

Materials, detailed methods, supplementary figures

## Acknowledgemets

We thank Behzad Rad for assistance with AFM imaging and Michael Connolly for consultation on peptoid synthesis. M.P.L. acknowledges support of a Burroughs Wellcome Fund Career Award at the Scientific Interface (CASI), a Stanley Fahn PDF Junior Faculty Grant with Award # PF-JFA-1760, a Beckman Foundation Young Investigator Award, a DARPA Young Faculty Award, an FFAR New Innovator Award, and a USDA award. M. P. L. is a Chan Zuckerberg Biohub Investigator and an Innovative Genomics Institute Investigator. L. C. is supported by National Defense Science and Engineering Graduate (NDSEG) Fellowship and by Lam Research. J. T. D. O. is supported by the Department of Defense office of the Congressionally Directed Medical Research Programs (CDMRP) Parkinson’s Research Program (PRP) Early Investigator Award. Work at the Molecular Foundry was supported by the Office of Science, Office of Basic Energy Sciences, of the U.S. Department of Energy under Contract No. DE-AC02-05CH11231, and the DARPA Fold F(x) program.

