## Supplemental Information for "Electrostatic-assemblies of single-walled carbon nanotubes and sequence-tunable peptoid polymers detect a lectin protein and its target sugars"

### Materials and Methods

**Materials** Reagents were purchased from Sigma-Aldrich, unless otherwise noted. Peptoid oligomers were synthesized on a Symphony X Synthesizer using the solid-phase submonomer method and purified by preparative reverse-phase HPLC, as previously described.[19] Beta-alanine tert-butylester hydrochloride was purchased from Chem-Impex International. 2-(2-(2-methoxyethoxy)ethoxy)-ethylamine was purchased from Peptide Solutions, Inc. All reagents were used without further purification. NeutrAvidin protein and Gibco Dulbecco's Modified Eagle Medium was purchased from Thermo Fisher Scientific. Trypsin gold was purchased from Promega. Proteinase K was purchased from New England Biosciences.

#### FRET binding assay for detecting WGA-peptoid interactions

The interactions between WGA and the L016 WGA-binding peptoid was validated using a FRET assay as previously described.[25] Briefly, nanosheet-forming peptoids with L016 loops were placed solution with 5 μM BODIPY-FL C16 and self- assembled by a rocking machine (rocking number was ~250 and the waiting time per rock was 10 s). Subsequently, the BODIPY-FL C16 incorporated peptoid nanosheets were mixed with the Alexa647-conjugated protein solutions at the following final concentrations: 10 mM nanosheets, 2.5 mM BODIPY-FL C16, and 250 nM Alexa-conjugated proteins in 25 mM phosphate buffer, 150 mM NaCl, pH 7.4.

#### Peptoid-SWNT adsorption

Adsorption of SWNT with peptoid was achieved using a protocol previously described.[44] Briefly, 1 mg of SWNT was added to 500 μL of phosphate buffered saline pH 7.4 and 100 nmol of peptoid. The solution was bath sonicated for 10 minutes, and then probe-tip sonicated using a Cole Parmer ultrasonic processor with pulses of 3-7 watts every 3 seconds for 5 minutes. The solution was subsequently allowed to equilibrate at the bench for 1 hour before centrifugation at 16.1 x $10^3$ Relative Centrifugal Force (RCF) for 30 minutes to remove any unsuspended nanotube aggregates. Assemblies were dark gray in color and concentration was characterized by UV-Vis-IR absorbance using a Shimadzu UV-3600 Plus. SWNT concentration is calculated from absorbance at 632 nm using Beer-Lambert law with extinction coefficient, $\varepsilon_{632} = 0.036$ L $mg^{-1}$ $cm^{-1}$.[7]

#### Near-infrared spectroscopy of peptoid-SWNT assemblies

All peptoid-SWNT nanosensor solutions were diluted to a final SWNT concentration of 5 mg/L in PBS. Spectroscopic analysis was performed by measuring the resulting SWNT photoluminescence with a home-built near infrared fluorescence microscope. Briefly, a Zeiss AxioVision inverted microscope was coupled to a Princeton Instruments IsoPlane 320 containing a liquid nitrogen-cooled Princeton Instruments PyLoN-IR 1D InGaAs array. The spectra of peptoid-SWNT samples were acquired, each in a separate well of a glass-bottom 384-well plate (Corning). Peptoid-SWNT assemblies were illuminated by a 500 mW, 721 nm laser reaching a sample equilibrium temperature of 37 ºC and a final power of 77 mW at the sample plane (SI 4a). All fluorescence spectral analysis was conducted using the (7,6) chirality emission peak, which is in high abundance in our SWNT samples.

**Proteolysis tests with peptoid-SWNT**

Peptoid-SWNT assemblies were diluted to a final concentration of 5 mg/L. Trypsin or proteinase K was added to the peptoid-SWNT assembly solutions to a final concentration of 1 mg/mL and 2 mg/mL, respectively. The solutions were incubated at room temperature and NIR spectra were collected 3 hours and 24 hours following the addition of protease. The fluorescence spectra were compared with a control where only buffer was added to the peptoid-SWNT sample.

**Protein screening experiments with peptoid-SWNT nanosensors**

Peptoid-SWNT nanosensors were diluted to a final working concentration of 5 mg/L. 27 μL of nanosensors were pipetted into each well of a 384-well glass-bottom plate and initial fluorescence spectra were recorded using NIR spectroscopy as detailed above. 3 μL of protein was added by pipetting to final protein concentrations as denoted in the manuscript, and fluorescence spectra were taken every 10 minutes following the addition of protein.

**Near-infrared microscopy of peptoid-SWNT nanosensors**

Peptoid-SWNT nanosensors are immobilized on a Mattek microwell dish with a 1.5 coverslip. The 1.5 coverslip surface was treated with 3-aminopropyltriethoxysilane (APTES) diluted to 10% (m/v) APTES in ethanol to create a positively charged surface onto which the nanosensors could immobilize. 100 μL of APTES solution was incubated on the coverslip for 2 minutes and followed by 3 washes with 1x PBS. 100 μL of peptoid-SWNT (20 mg/L) was incubated on the surface for 5 minutes, and followed by 3 washes with 1x PBS to rid the surface of unbound nanosensors. Single peptoid-SWNT nanosensors were excited with a 500 mW, 721 nm laser with a final power of 77 mW at the objective, and imaged using a ZEISS α Plan-APOCHROMAT 100x oil immersion objective (numerical aperture (NA) = 1.46) on a Zeiss AxioVision inverted microscope coupled to a Princeton Instruments NIRVana 640 InGaAs camera. Regions of interest were identified around areas where initial fluorescence was over 5-fold the fluorescence of the background.

**Detection of wheat germ agglutinin protein in cell medium**

Peptoid-SWNT nanosensors were diluted in Gibco Dulbecco's Modified Eagle Medium to a SWNT concentration of 5 mg/L. The nanosensors were allowed to equilibrate for 2 hours before spectroscopy using the NIR spectrometer as described above. WGA was added to the well to a final concentration of 10 µM.

**Detection of sugars using peptoid-SWNT nanosensors**

Peptoid-SWNT nanosensors were diluted to a SWNT concentration of 5 mg/L in PBS. Wheat germ agglutinin protein was added to the nanosensors to a final concentration of 10 µM, and the nanosensors were allowed to equilibrate for 1.5 hours before spectroscopy using the NIR spectrometer described above. 1 mM final concentration of sugar was added to each well and spectra were acquired every 10 minutes for 1 hour.

**Surfactant-induced solvatochromic shift of peptoid-SWNTs**

Peptoid-SWNT nanosensors were diluted to a SWNT concentration of 5 mg/L in PBS and imaged with NIR fluorescence spectroscopy as detailed above. NIR fluorescence spectra of 27 μL peptoid-SWNTs were acquired every 10 minutes for 30 minutes before and after the addition of 3 μL 5% (w/v) sodium cholate for a final 0.5% (w/v) concentration of sodium cholate.

**Temperature Equilibration Experiments**

NIR fluorescence measurements of peptoid-SWNTS were performed with illumination by a 721 nm, 500 mW laser. We measured the temperature of a 30 μL sample well in a 384-well Corning glass bottom plate with continuous laser illumination over 60 minutes, representing the standard time-course of our peptoid-SWNT fluorescence assays, using a FLIR thermal camera. Temperature measurements were taken every 10 minutes, with additional timepoints taken at 30 seconds and 1 minute.

**Atomic Force Microscopy of Peptoid-SWNT Nanosensors**

Monodispersed peptoid-SWNT nanosensors were analyzed with atomic force microscopy using an Asylum Research MFP-3D AFM. 20 μL of peptoid-SWNT assemblies (20 mg/L) in PBS were deposited on freshly cleaved mica, and incubated for 1 hour at room temperature. Unbound nanosensors and salts were washed from the mica 3 times using MilliQ water. For AFM of protein on nanosensors, ProLoop-SWNT nanosensors (20 mg/L) were incubated with 10 μM WGA for 1 hour and the samples were next deposited on freshly cleaved mica and incubated for an additional hour. Unbound nanosensors, salts, and proteins were washed 3x with MilliQ water prior to AFM imaging.

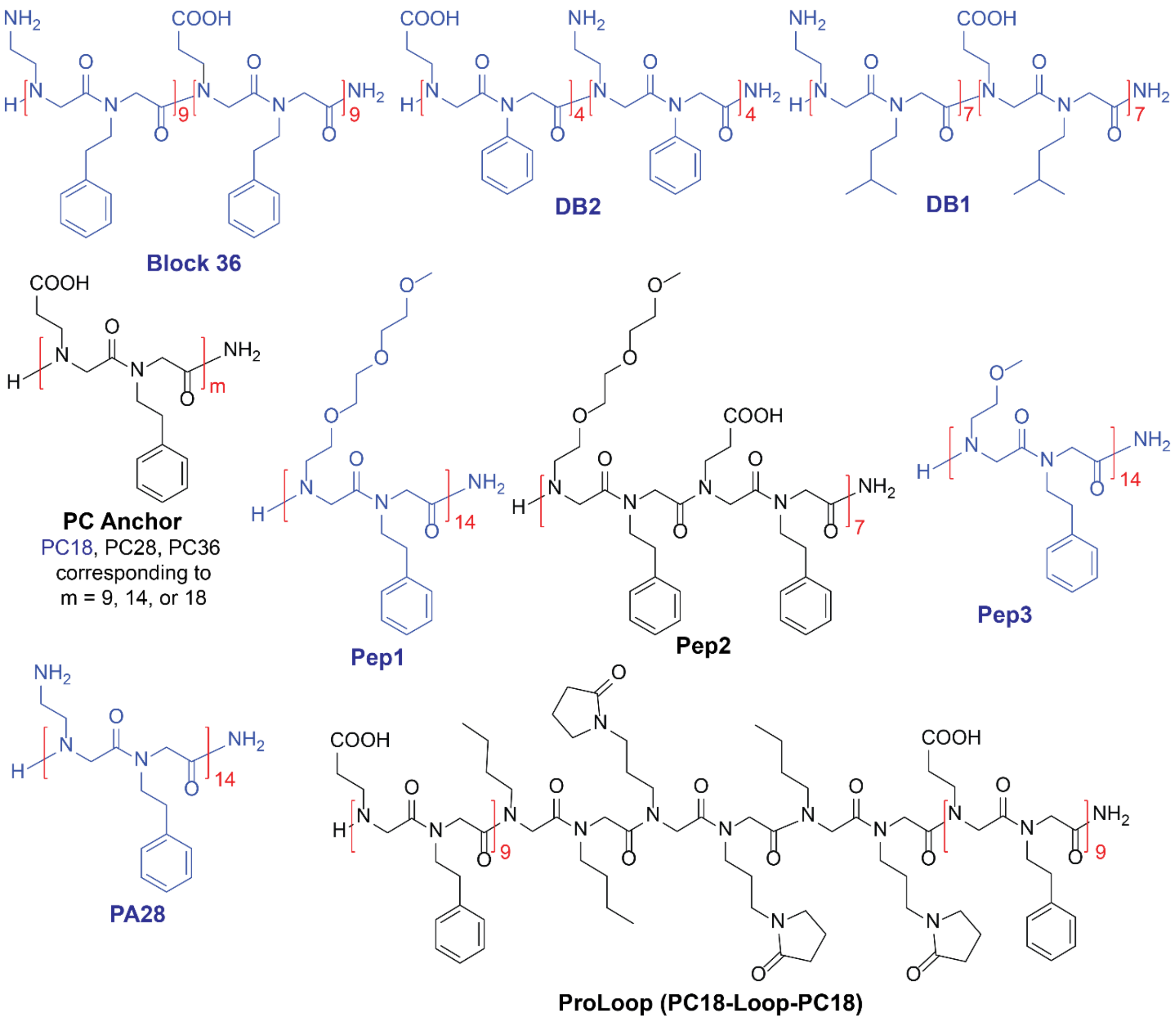


**SI. 1: Peptoid Structures.** Expanded chemical structures of the 11 peptoid polymers in our library. Blue peptoids denote sequences that did not adsorb to SWNT, and black peptoids denote sequences that successfully adsorbed to form peptoid-SWNT assemblies.

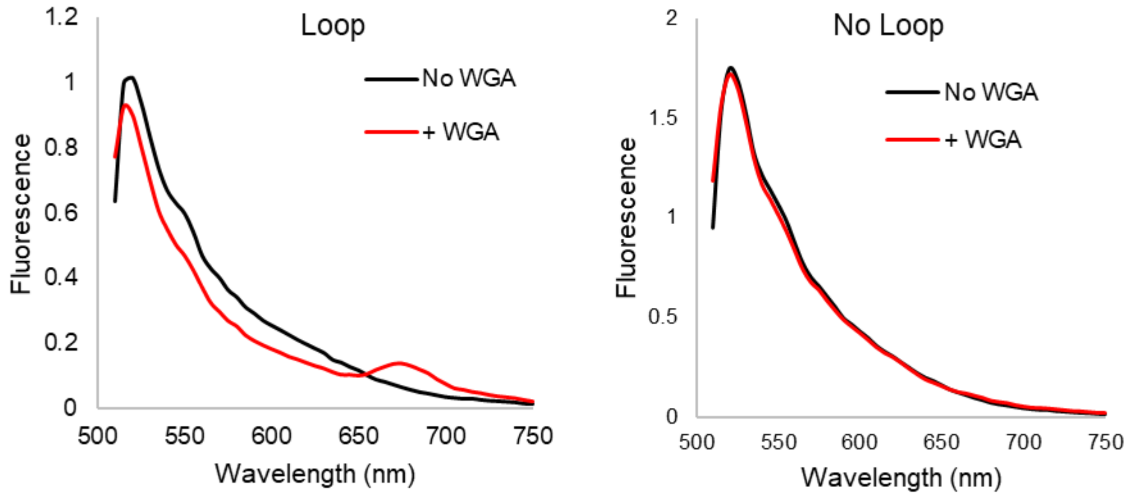


**SI. 2: FRET Assay Showing Peptoid Loop Interaction with WGA**. A FRET assay shows interaction between the 6-monomer peptoid loop and WGA protein. Peptoid nanosheets with and without peptoid loops were assembled with BODIPY-FL C16 dye (FRET donor) with a peak emission wavelength at 528 nm. The addition of Alexa647-labeled WGA (FRET acceptor) to peptoid-containing loops exhibited in a decrease in donor fluorescence and an increase in acceptor fluorescence indicating an increase in FRET. The FRET ratio (calculated as the ratio of increase in acceptor fluorescence to decrease in donor fluorescence) for peptoid with the peptoid loop was 0.678, while the FRET ratio for peptoid without loops was -0.172.

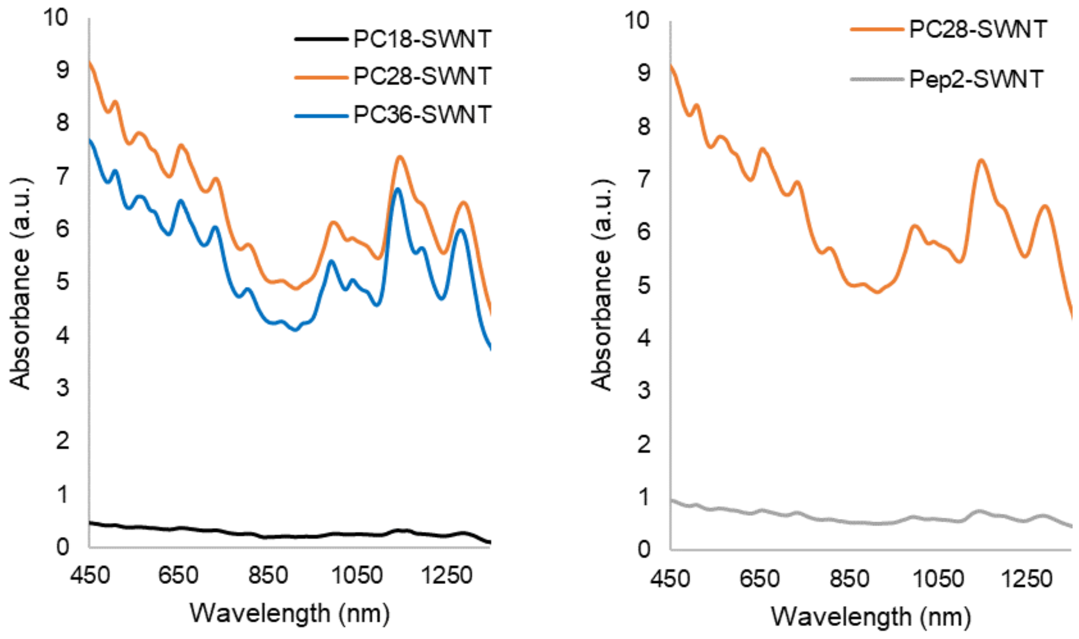


**SI. 3. Absorbance Spectra of SWNT.** Peptoid suspension efficiency to SWNTs was measured by the optical absorption of the sample at 632 nm, from a broad-spectrum absorption scan by a UV-Vis-IR spectrophotometer. SWNT concentration is calculated from absorbance at 632 nm using the Beer-Lambert law with extinction coefficient, $\varepsilon_{632} = 0.036$ L mg$^{-1}$ cm$^{-1}$.[1]

**SI. 4. Characterization of Peptoids - Analytical High Performance Liquid Chromatography Spectra**

HPLC spectrum for PC18

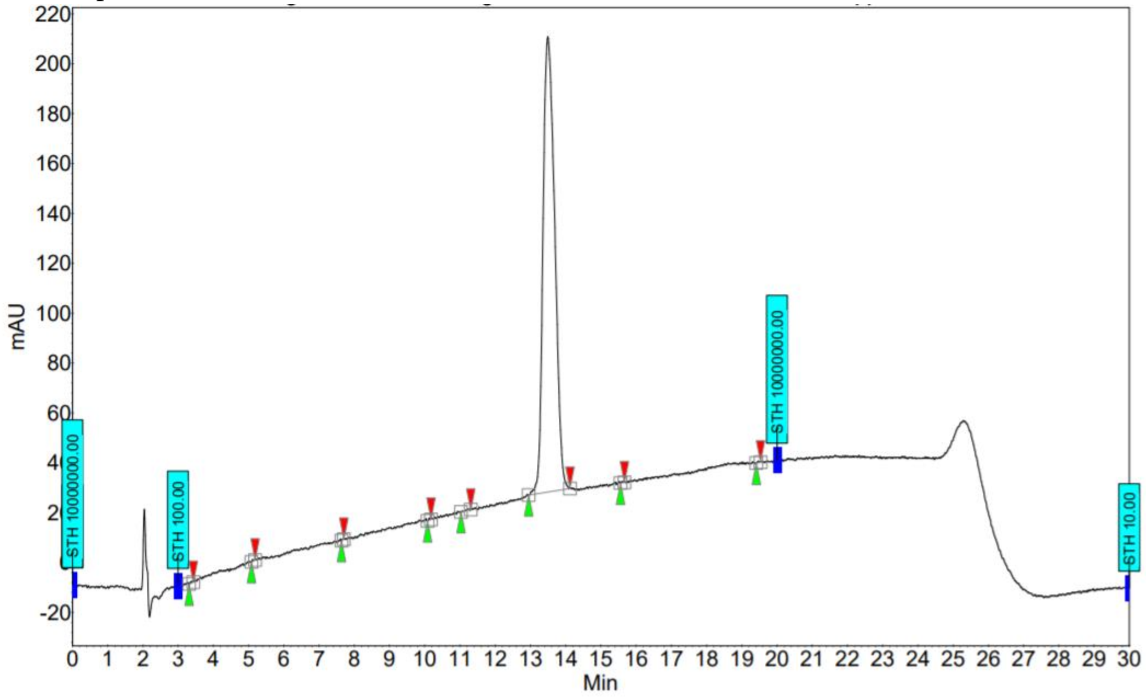


HPLC spectrum for PC28

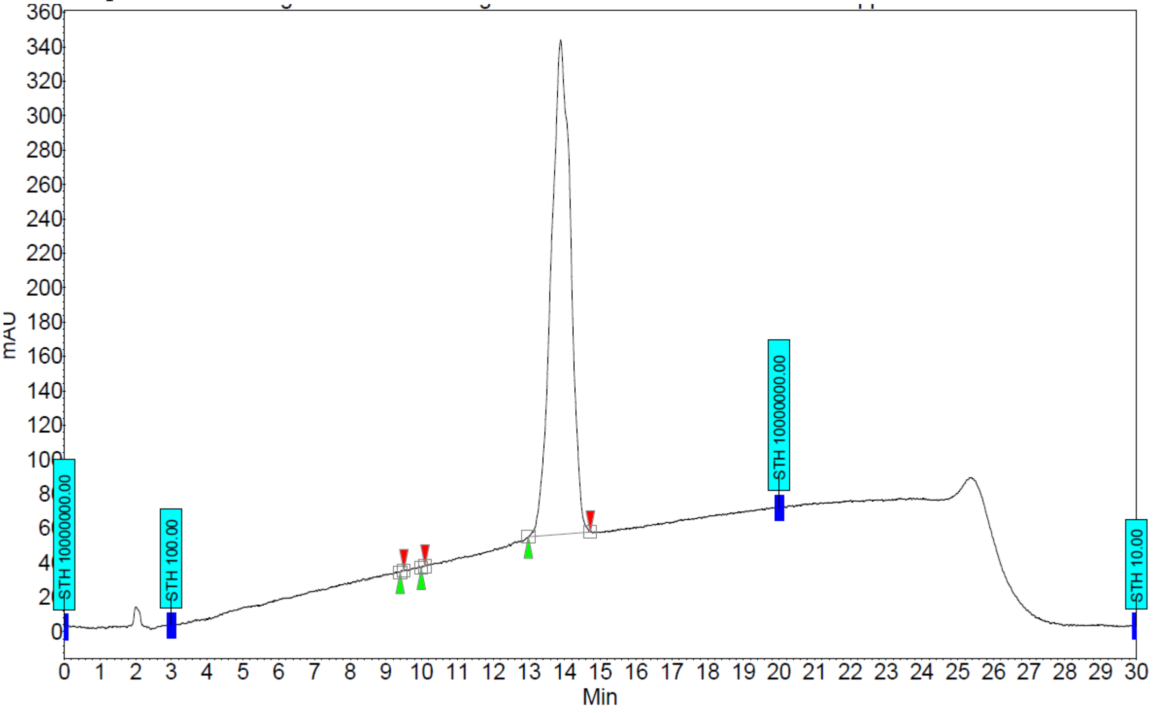

HPLC spectrum for PC36

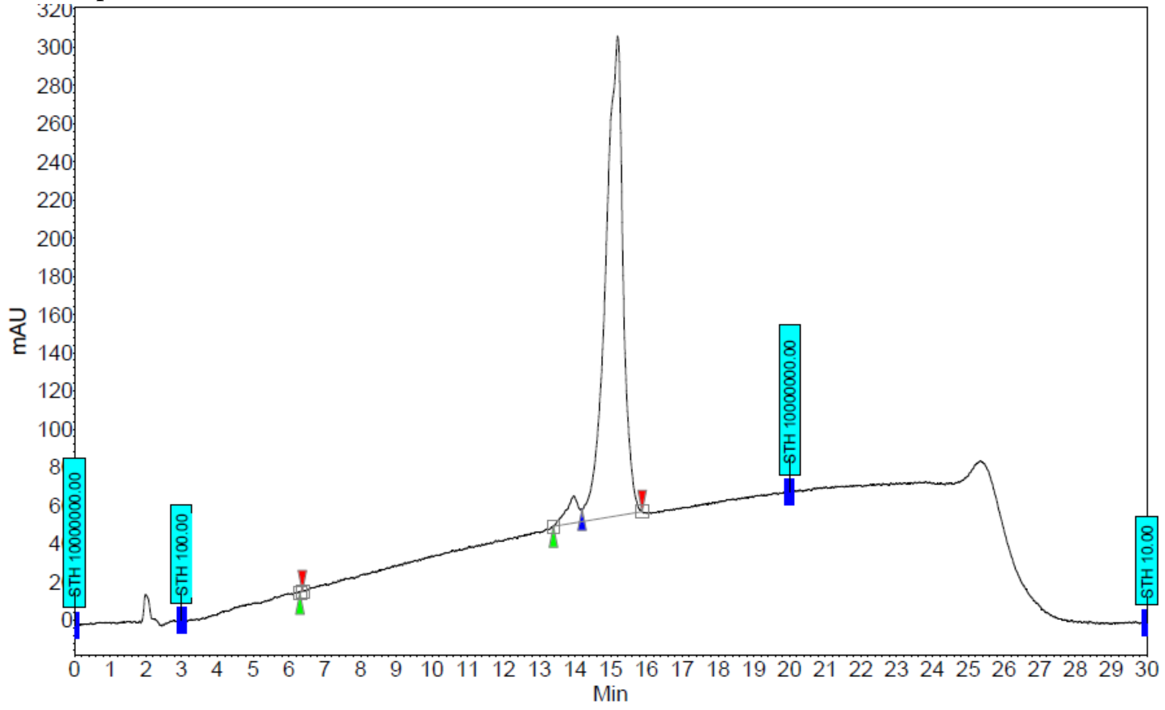


HPLC spectrum for ProLoop

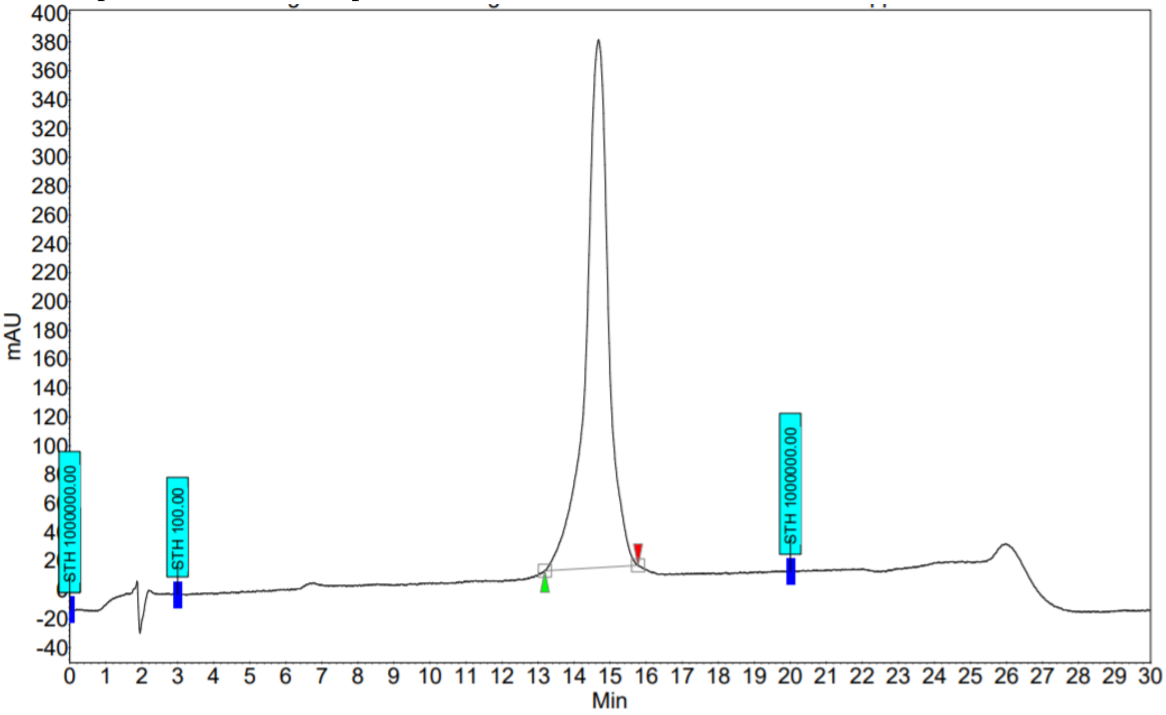

HPLC spectrum for Pep1

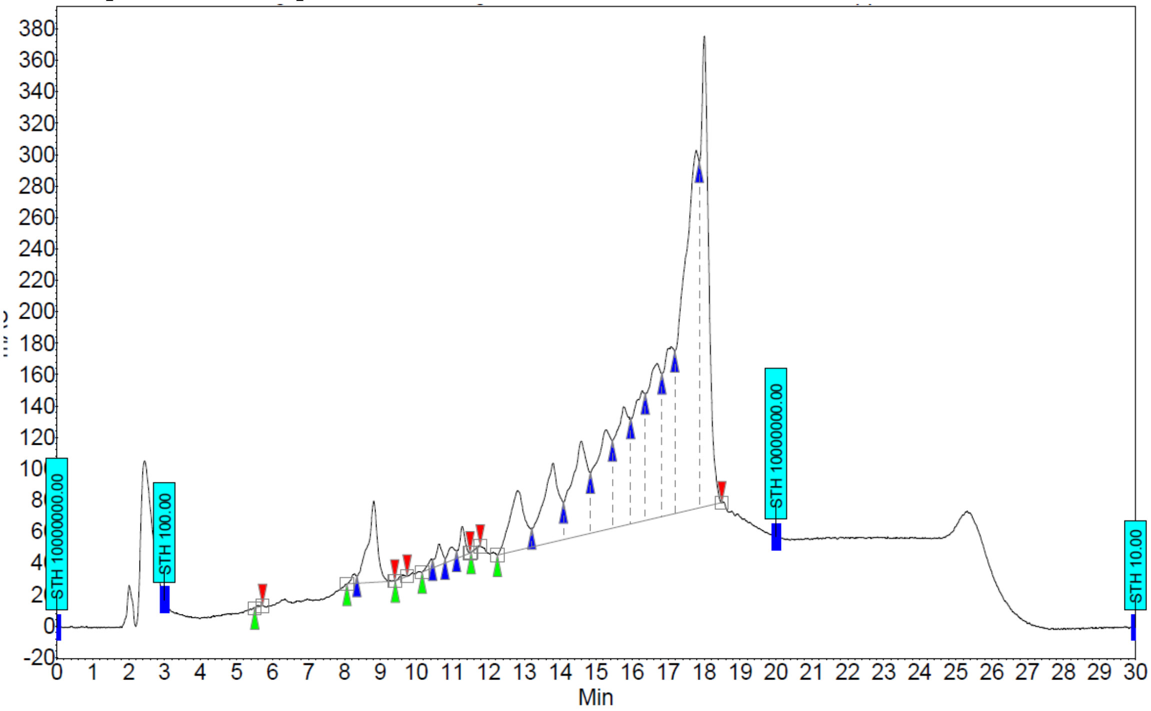


HPLC spectrum for Pep2

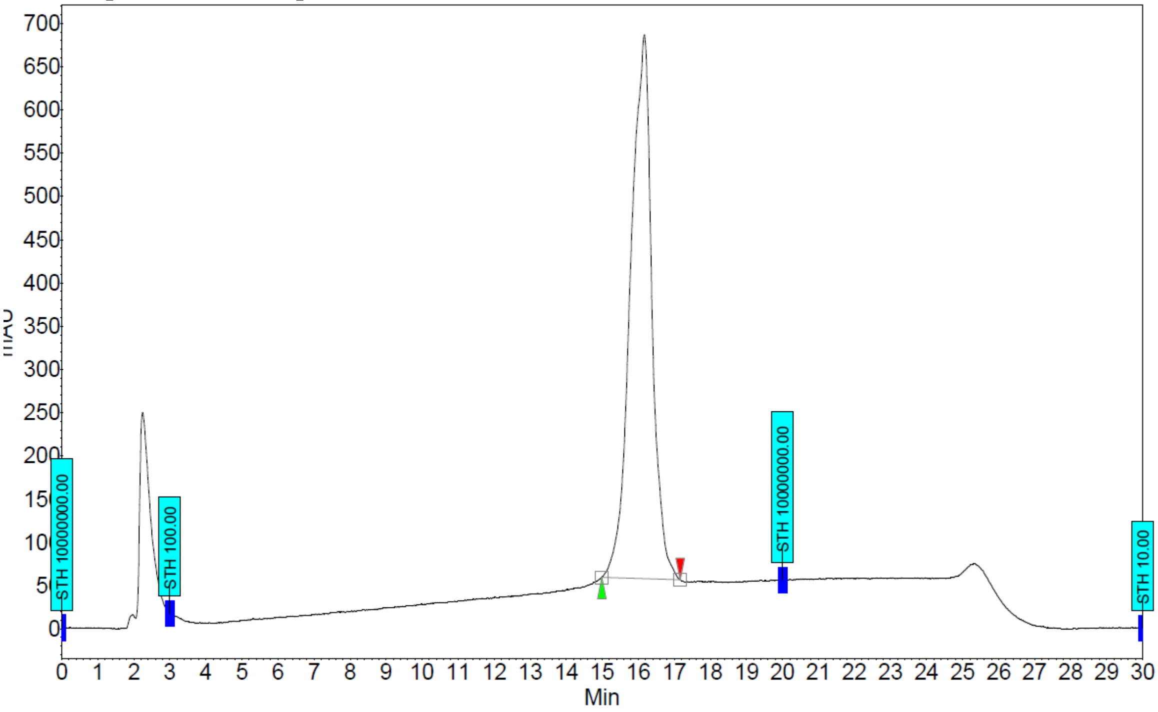

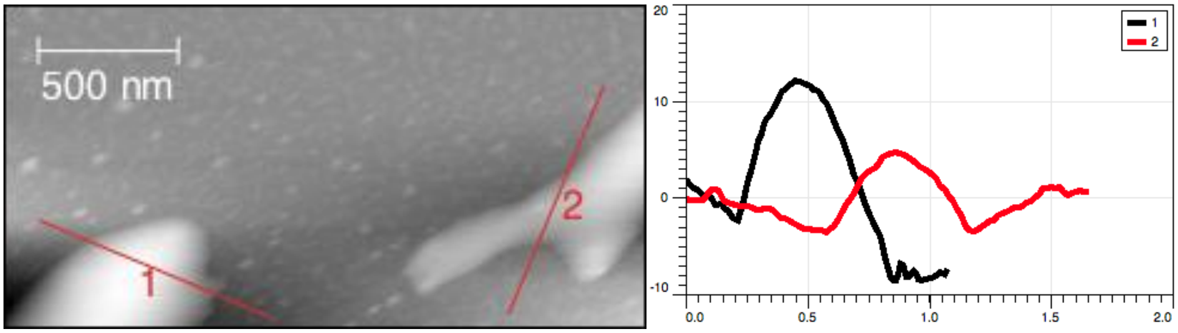


**SI. 5. Block 36 Instability in Peptoid-Suspension.** Block36 (B36) peptoid has been previously characterized and shown to form self-assembled nanosheets that are absent in PA28 and PC28.[2] Similarly, following probe-tip sonication with SWNT, B36 peptoids yielded unstable colloidal SWNT suspensions and large aggregates (10 nm in height) were observed in AFM, suggesting B36 peptoid nanosheet formation is thermodynamically favorable over B36-SWNT assembly formation. x/y axes for traces 1 and 2 are length in nm.

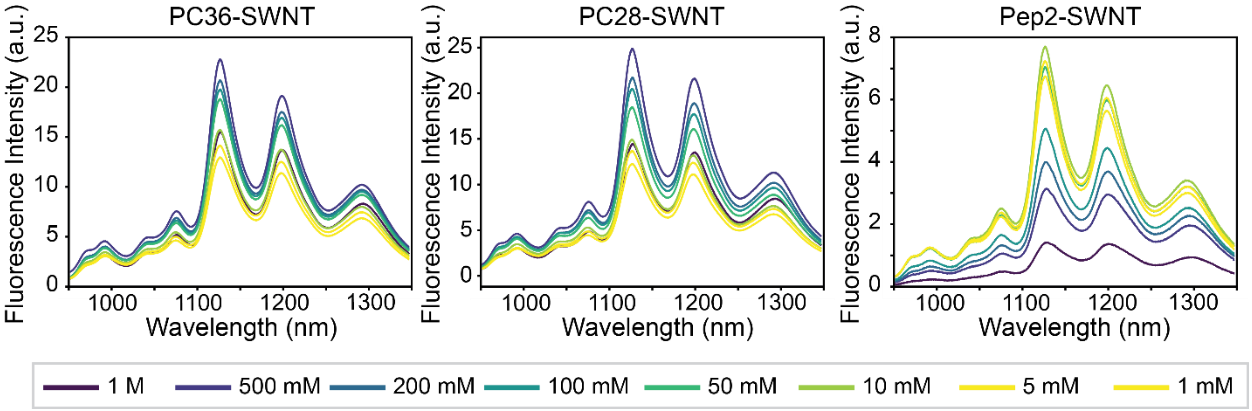


**SI. 6 Peptoid-SWNT Fluorescence Under Variable Ionic Strength Conditions.** Solution ionic strength affects the fluorescence of peptoid-SWNT assemblies, whereby higher ionic strengths yield higher peptoid-SWNT fluorescence. Pep2-SWNT exhibits low fluorescence intensities for all salt concentrations greater than 50 mM due to colloidal instability and aggregation.

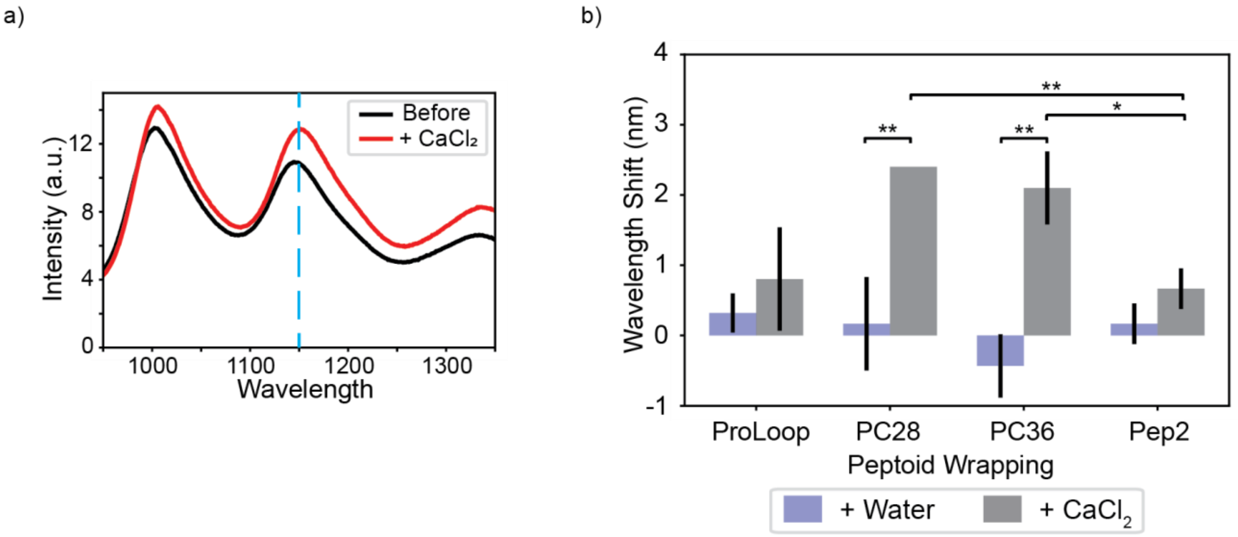


**SI. 7. Divalent Salt Experiments to Test Peptoid-SWNT Conformation.** a) Peptoid-SWNT assemblies, such as PC36-SWNT, exhibit a solvatochromic shift upon addition of divalent cations from $CaCl_2$. b) Comparison of the solvatochromic shift at the 1195 nm peak shows a significant shift upon addition of $CaCl_2$ to PC28-SWNT and PC36-SWNT as compared to water addition, although all peptoid-SWNT showed a red-shift in fluorescence wavelength upon addition of 500 mM $CaCl_2$. Notably, PC28-SWNT and PC36-SWNT, which have the same monomeric composition, show a significantly different response to divalent salts compared to Pep2-SWNT (* denotes $p < 0.05$, ** denotes $p < 0.005$).

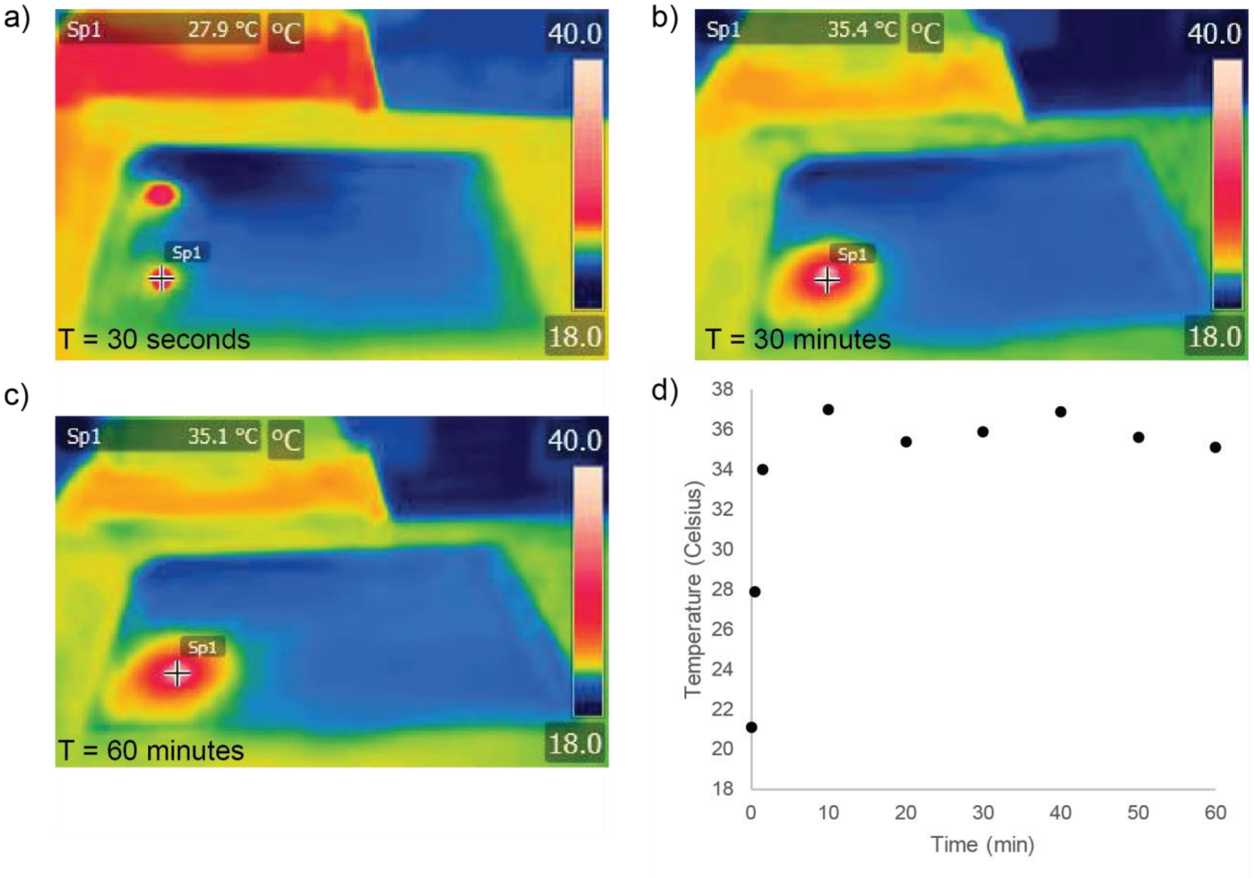


**SI. 8. Temperature Equilibration of Peptoid-SWNT Fluorescence Spectral Assays.** FLIR thermal camera images of the sample well (Sp1) at a) 30 seconds, b) 30 minutes, and c) 60 minutes. d) Time-dependent temperature changes show rapid temperature equilibration in the well with a maximum temperature stabilizing at ~37 ºC.

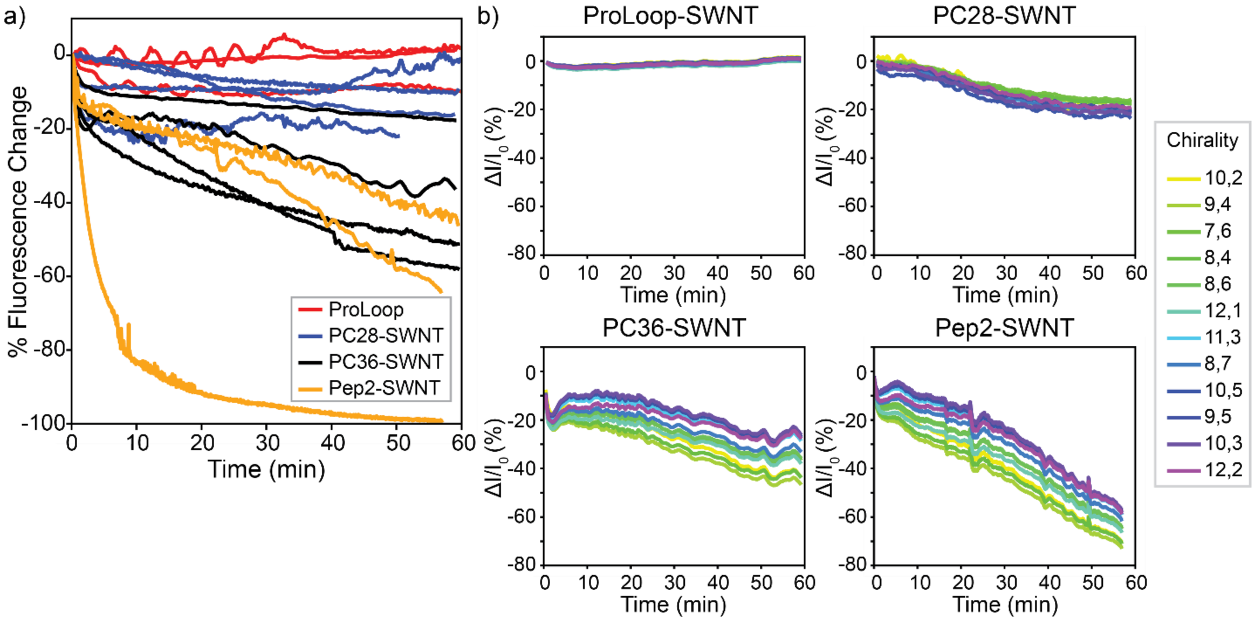


**SI. 9: Peptoid-SWNT Thermostability Tests.** a) Plot of all trials performed to determine peptoid-SWNT assemblies' thermostability. ProLoop-SWNT is the most stable peptoid-SWNT assembly under continuous laser illumination, whereas Pep2-SWNT is the least stable. b) Chirality-independent changes in fluorescence to heating.

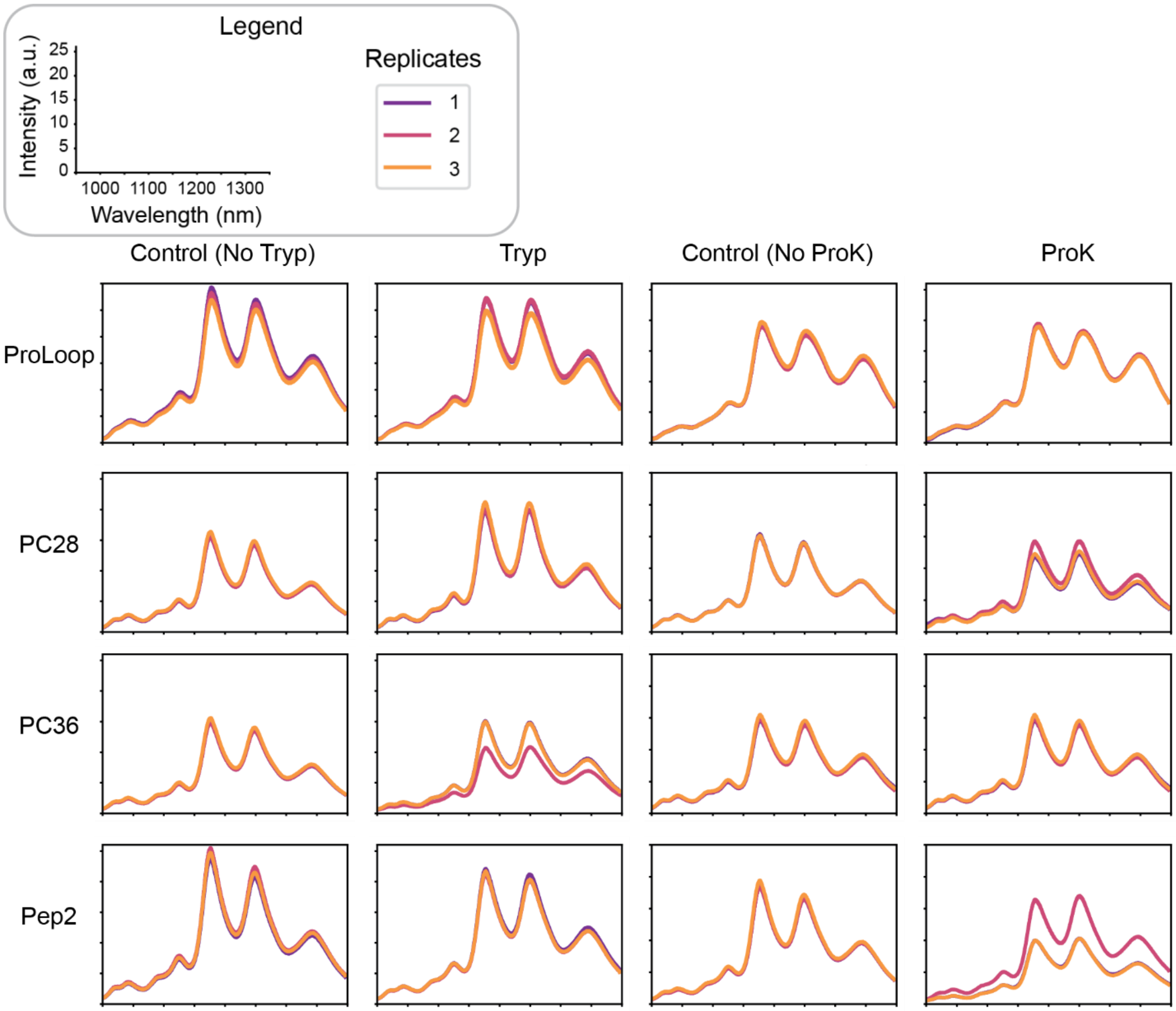


**SI. 10. Protease Exposure Demonstrates Peptoid-SWNT Stability.** Fluorescence spectra of peptoid-SWNT assemblies following 24 hours of room temperature incubation with trypsin (Tryp) or Proteinase K (ProK). A decrease in fluorescence of the sample with protease with respect to a control sample containing no protease suggests protease-induced aggregation. All peptoid-SWNTs retained fluorescence upon exposure to proteases, with ProLoop-SWNT exhibiting the greatest stability.

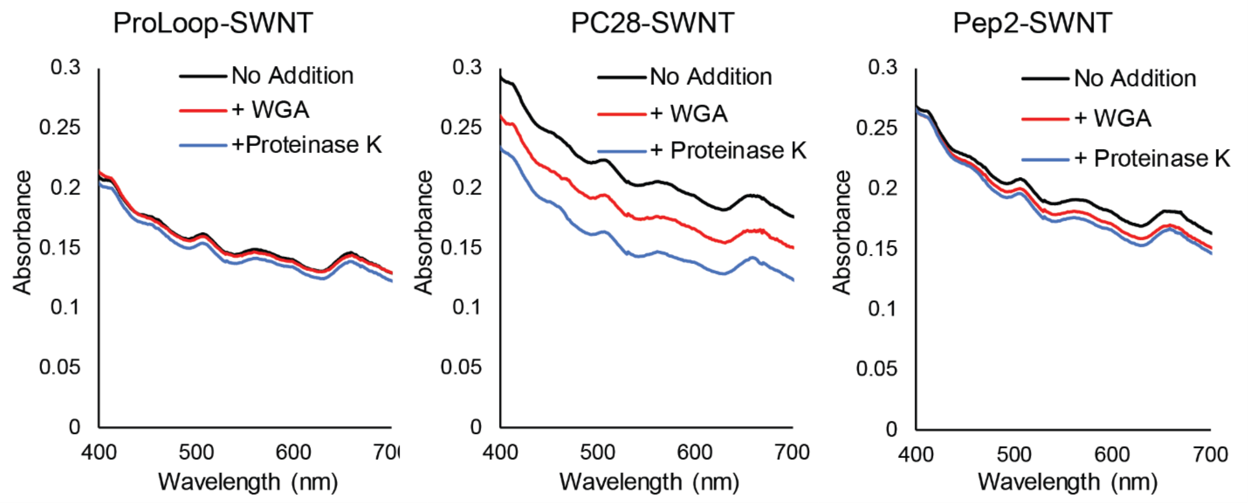


**SI. 11. Assessing Peptoid-SWNT Aggregation.** Aggregation of peptoid-SWNT assemblies in the presence of WGA or Proteinase K was assessed using a protein turbidity assay.[3] Peptoid-SWNT conjugates were incubated with either protease for 24 hours prior to acquiring the sample UV-Vis-IR absorbance spectra. ProLoop-SWNT shows no loss of absorbance following incubation with WGA and minimal loss of absorbance following incubation with Proteinase K suggesting ProLoop-SWNT stability.

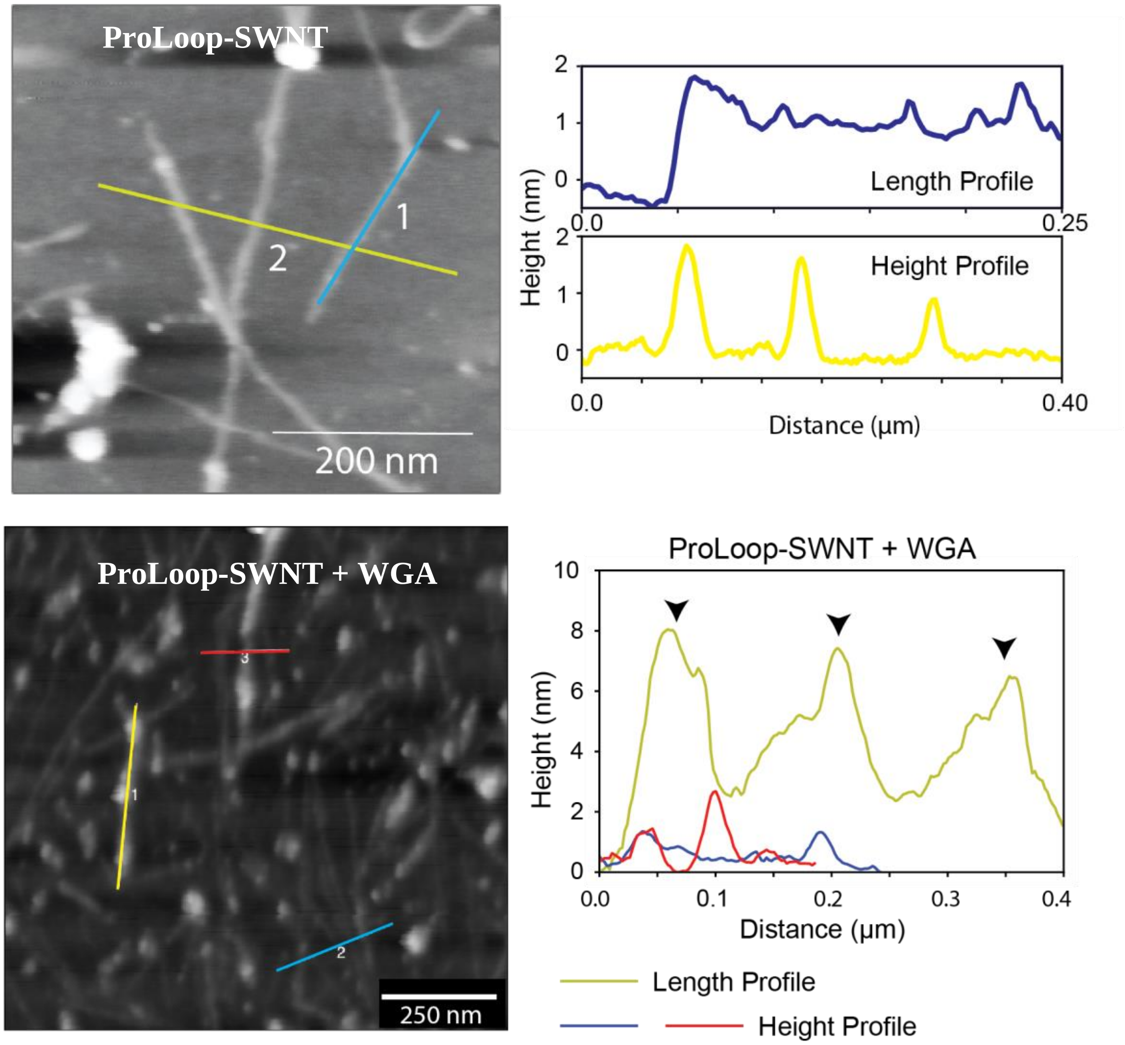


**SI. 12. AFM of Peptoid-SWNT and Peptoid-SWNT + WGA Protein.** AFM imaging of ProLoop-SWNT nanosensors shows heights of 1 nm to 2 nm, and ProLoop-SWNT + WGA heights of 6 nm to 8 nm, which is expected from adsorption of a 36 kDa, or ~2 nm globular protein such as WGA. Several globular features 6-8 nm in size were observed along the length of the ProLoop-SWNT following incubation with WGA.

**SI 13. Curve Fitting to Determine Goodness of Fit of Equilibrium Model**

To determine the limit of detection of ProLoop-SWNT nanosensor to WGA, the change in nanosensor fluorescence was fit to a model of enzyme to substrate binding. An extended range of the WGA concentrations for nanosensor response is found below. Inset: ProLoop-SWNT response to 0 μM to 1 μM WGA.

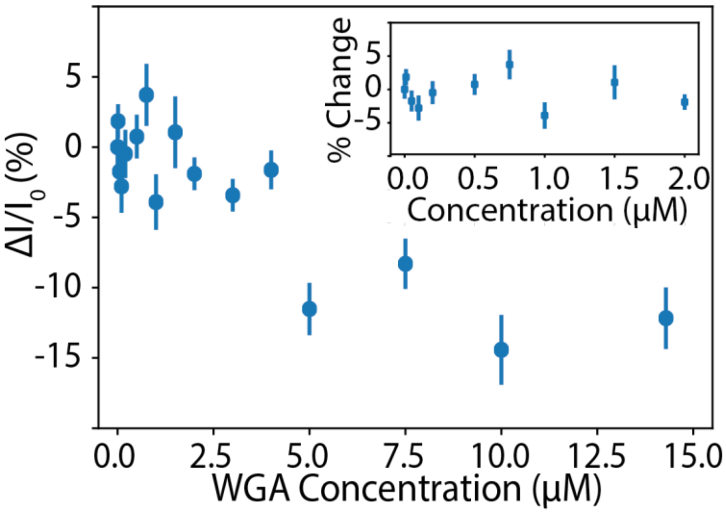


Assuming each fluorescent site binds one WGA protein, the fluorescence of the nanosensor can be treated as directly proportional to the number of protein-bound sites on the nanosensor.
The protein binding sites are assumed to form a monolayer, with each nanosensor site binding one protein site, thus WGA binding is described by:

$$WGA + \theta \leftrightarrow [WGA\text{-}\ \theta]$$

At equilibrium the equilibrium constant is given as:

$$K = \frac{[WGA-\theta]}{[WGA][\theta]}$$

The total concentration of available binding sites is a constant value $[\theta]_{tot}$ given by the sum of the free [θ] and protein-bound states [WGA- θ]:

$$[\theta]tot = [\theta] + [WGA\text{-}\ \theta]$$

$$= \frac{[WGA-\theta]}{K[WGA]} + [WGA\text{-}\ \theta]$$

$$= [WGA - \theta]\left(1 + \frac{1}{[\mathrm{WGA}]K}\right)$$

$$= [WGA - \theta]\left(\frac{[\mathrm{WGA}]K+1}{[\mathrm{WGA}]K}\right)$$

Assuming the nanosensor intensity change is proportional to the number of bound sites out of the total available nanosensor sites:

$$\frac{I - I_0}{I_0} = a\frac{[WGA - \theta]}{[\theta]_{tot}} \; = \; a\frac{(K[WGA])^n}{(K[WGA])^n + 1} + b$$

Where n accounts for WGA binding cooperativity. Since WGA is functional as a dimer, we restrict n to be a maximum of 2 for the fit to converge with WGA binding data.

We plotted residual values for the goodness of fit of the above model, for which we obtained a scattered distribution of residual values (uncorrelated residuals), suggesting the model is not inherently biased by the data.

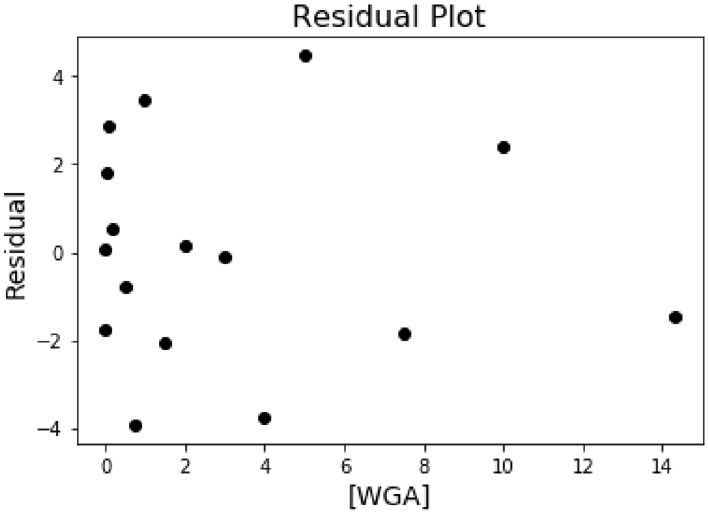


The sum of residuals given by an n = 2 fit is $\Sigma_{res}$ = 0.0164.

To calculate the limit of detection (LoD) of the ProLoop-SWNT nanosensor, we defined the LoD as the lowest analyte concentration likely to be reliably distinguished from the limit of blank (LoB).[4] The limit of blank is the highest apparent analyte concentration expected to be found when replicates of a blank sample containing no analyte are tested.

LoB = $\text{mean}_{\text{blank}}$ + 1.645 ($\text{SE}_{\text{blank}}$)

LoD = LoB + 1.645 ($\text{SE}_{\text{low concentration sample}}$)
For our system, LoB = 0 + 1.645 (-1.3904)

LOD = LoB + 1.645 (-1.2107) = -4.2788 (fluorescence change value) which corresponds to a WGA concentration of 3.4 μM.

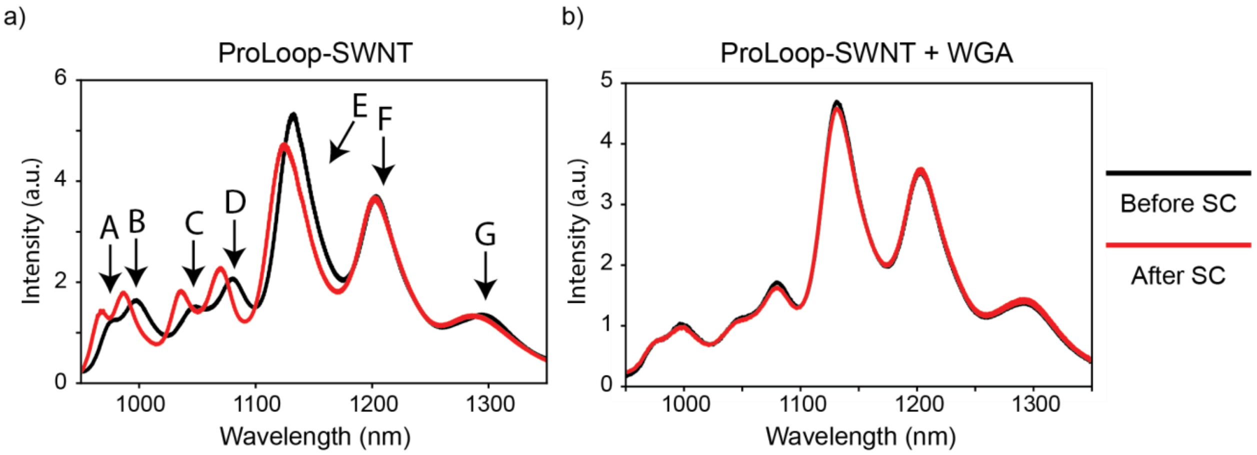


c)

| Peak | Initial Wavelength (nm) | Peak Shift (nm) | |
|---|---|---|---|
| | | - WGA | + WGA |
| A | 974.89 | -7.88 | 2.44 |
| B | 998.67 | -10.33 | 0.98 |
| C | 1045.47 | -11.76 | 0.0 |
| D | 1080.62 | -10.75 | 0.0 |
| E | 1130.50 | -7.30 | 0.0 |
| F | 1202.29 | -0.97 | 0.0 |
| G | 1295.06 | -7.67 | 0.95 |

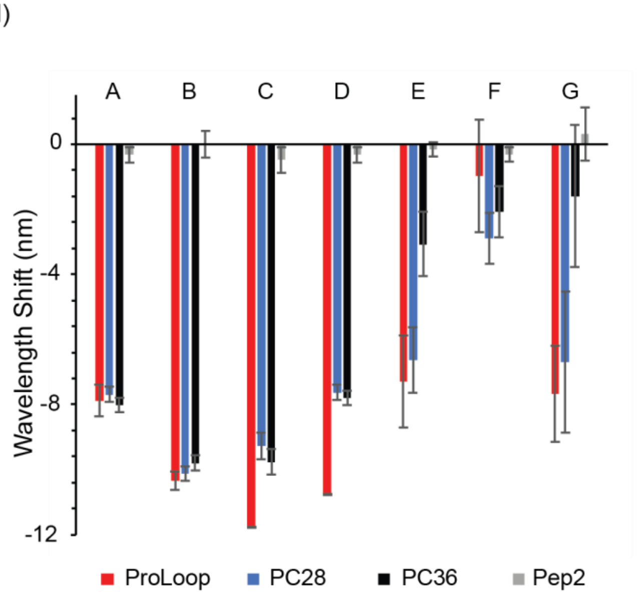


**SI. 14. ProLoop-SWNT Nanosensor Sodium Cholate Stability Tests.** a) Introduction of 0.5% (w/v) sodium cholate to the ProLoop-SWNT nanosensor induced solvatochromic shifts in the peptoid-SWNT nanosensors. b) Pre-incubation of ProLoop-SWNT nanosensors with WGA prior to introduction of 0.5% sodium cholate eliminated the solvatochromic shift, indicating WGA binds to and stabilizes the ProLoop-SWNT assembly. c) Peak wavelength shifts for multiple chiralities of ProLoop-SWNT. d) Peak wavelength shifts observed for ProLoop, PC28, PC36, and Pep2 peptoid-SWNT assemblies. Notably, Pep2-SWNT is invariant to the addition of SC.

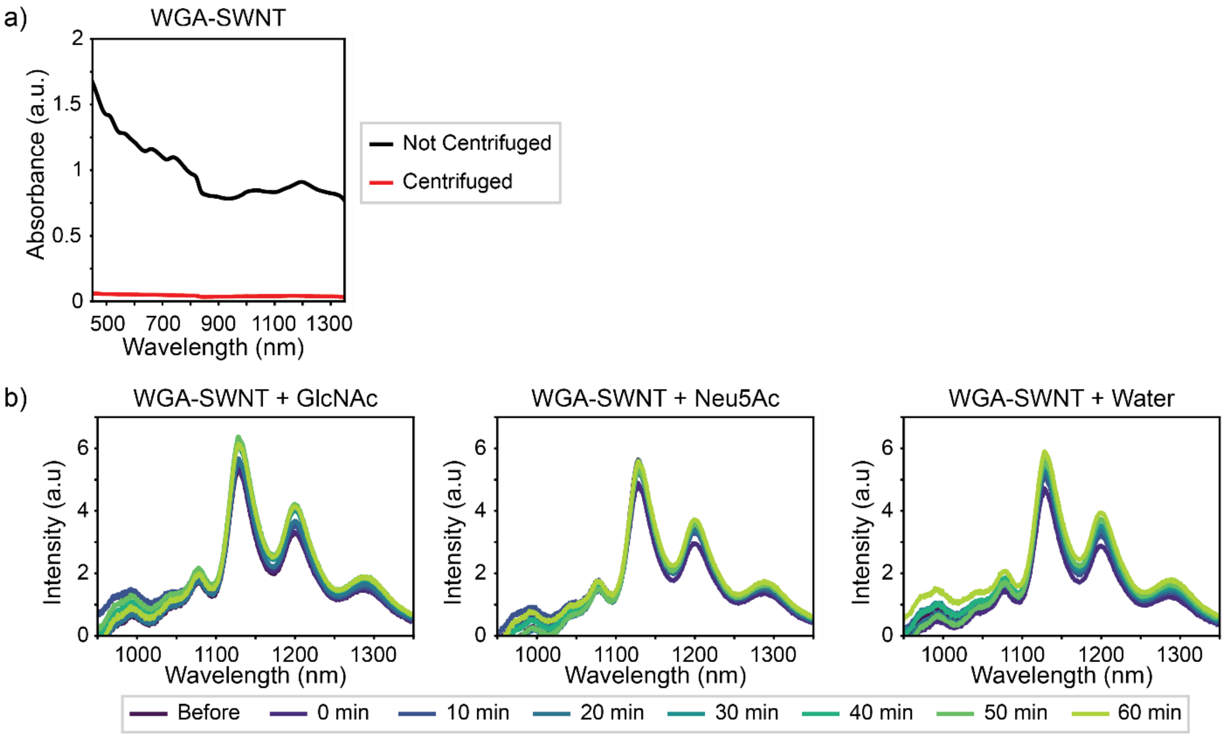


**SI. 15. WGA Interactions with SWNT.** a) Absorbance spectrum of SWNT sonicated with WGA. The WGA-SWNT complex was only loosely associated, and centrifugation of WGA-SWNT caused aggregation and subsequent the loss of absorbance and fluorescence. b) Representative fluorescence spectra of the WGA-SWNT suspension without centrifugation shows no response to the WGA target sugars GlcNAc and Neu5Ac.

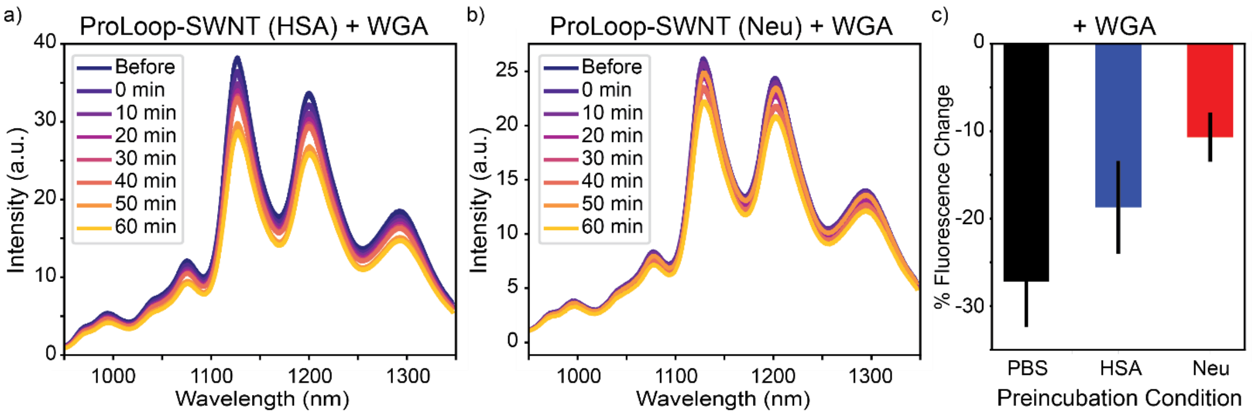


**SI. 16. ProLoop-SWNT Nanosensors Show Response to WGA in Protein-Rich Environments.** The response of ProLoop-SWNT nanosensors to WGA in protein-rich conditions; parentheses indicate the pre-incubation conditions of the ProLoop-SWNT nanosensor an hour before the start of the imaging experiment in a) human serum albumin (HSA, 40 mg/mL concentration) and b) NeutrAvidin (Neu, 5 mg/mL concentration). c) ProLoop-SWNT response to WGA following incubation with these proteins is attenuated as compared to ProLoop-SWNT nanosensors in PBS buffer indicating some sensing interference by external proteins. Error bars denote standard deviation.

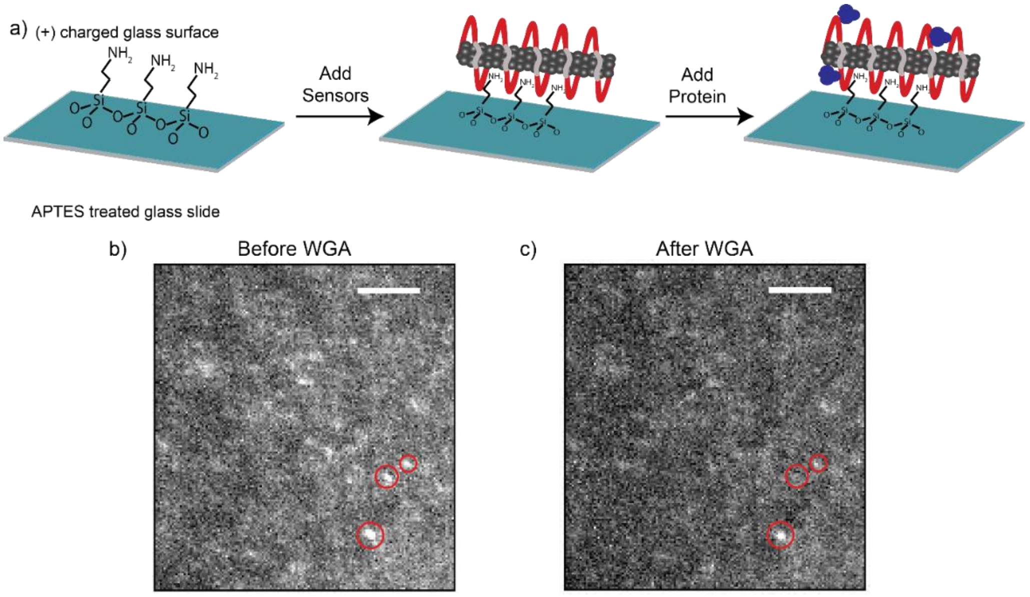


**SI. 17. NIR Microscopy of ProLoop-SWNT Nanosensors.** a) Schematic of single nanosensors immobilized on a (3-aminopropyl)triethoxysilane treated surface for NIR microscopy. b) Representative image of ProLoop-SWNT nanosensors seen in NIR microscopy, where red circles indicate regions of interest around nanosensors where fluorescence traces were recorded before and after the addition of WGA. c) Same field of view after addition of WGA that caused a loss of fluorescence to the nanosensors as discussed in Figure 4c. Scale bars indicate 25 µm.

### ProLoop-SWNT Sugar Sensing Controls

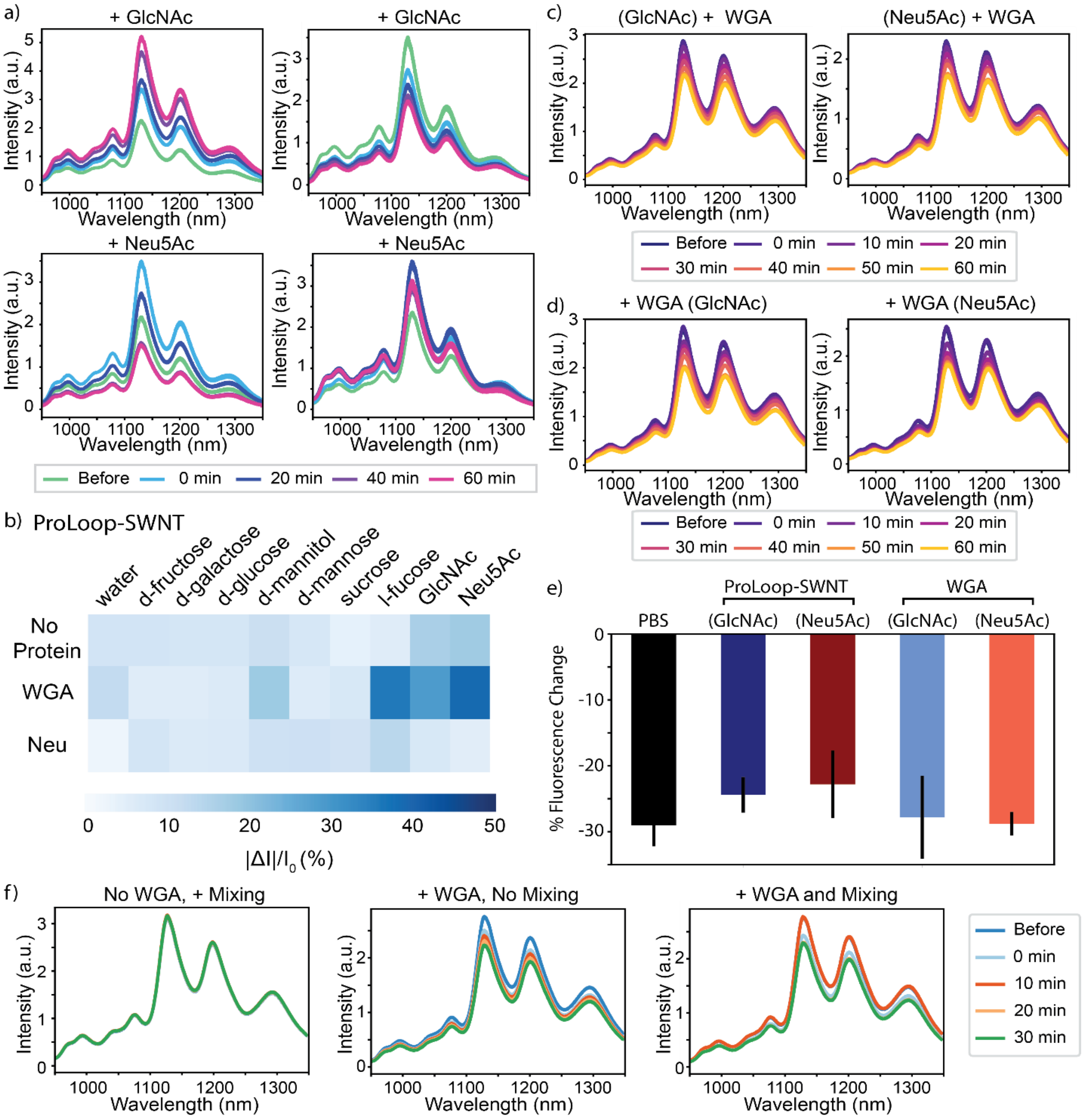


**SI. 18. Ternary Sugar Sensing Controls.** a) WGA-ProLoop-SWNT complexes non-monotonically respond to WGA's target sugars. b) WGA-ProLoop-SWNT nanosensors response to WGA's target sugars is not seen in the absence of WGA or the presence of NeutrAvidin (Neu) instead of WGA. c) Pre-incubation of ProLoop-SWNT with WGA's target sugars (parenthesis indicate pre-incubation conditions) did not impede nanosensor response to WGA. d) Pre-incubation of WGA with its target sugars (parenthesis indicate pre-incubation conditions) did not impede ProLoop-SWNT nanosensor response to WGA. e) Sugar blocking experiment summary (parenthesis indicate pre-incubation conditions of either ProLoop-SWNT or WGA). f) Mixing experiments showed that sample agitation is not the cause of nanosensor response to WGA.

ProLoop-SWNT nanosensors, following complexation with WGA, exhibit a fluorescence response to WGA's target sugars *N*-acetyleglucosamine (GlcNAc) and *N*-acetyleneuraminic acid (Neu5Ac). The fluorescence response of the ProLoop-SWNT nanosensor + WGA to sugar inducs a non-monotonic increase or decrease in fluorescence, and is quantified as the absolute value of intensity change (SI. 18a). ProLoop-SWNT nanosensors without WGA did not respond to the addition of target sugars (SI. 18b). Furthermore, ProLoop-SWNT pre-incubated with another protein, Neutravidin, also did not respond to the addition of target sugars. These results suggest specificity of the ProLoop-SWNT-WGA assembly towards its target sugars through WGA lectin recognition, and these signals are not attributed to the ProLoop-SWNT alone.

To assess whether target sugars affect the ability of ProLoop-SWNT nanosensors to detect WGA, we pre-incubated ProLoop-SWNT with target sugars GlcNAc and Neu5Ac an hour before measuring binding between WGA and ProLoop-SWNT (Fig. 18c). Pre-incubation with sugar did not hinder ProLoop-SWNT sensing of WGA, with the magnitude of fluorescence change upon the addition of WGA similar for each pre-incubation condition. Similarly, pre-incubation of the WGA protein with GlcNAc and Neu5Ac did not hinder ProLoop-SWNT sensing of WGA (Fig. 18d). These controls yielded similar magnitudes of fluorescence response of ProLoop-SWNT to WGA (SI. 18e).

We further showed the fluorescence modulation of the ProLoop-SWNT nanosensor towards WGA is not affected by or due to physical agitation (Fig. 18f). Mixing of WGA-ProLoop-SWNT complexes by pipetting did not induce WGA-ProLoop-SWNT fluorescence modulation in the absence of target sugars.

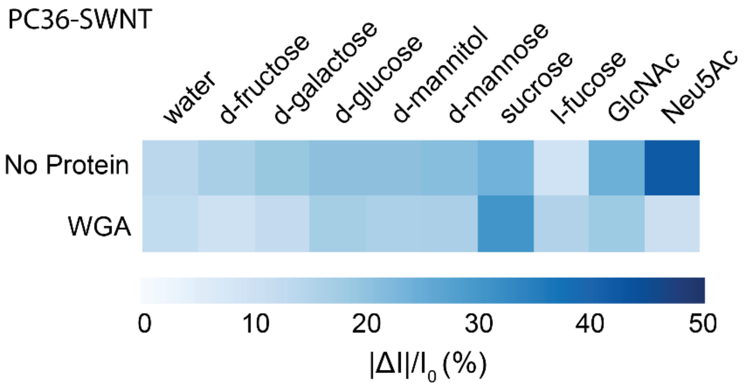


**SI. 19. PC36 Peptoid-SWNT + WGA Response to Sugar Panel.** PC36-SWNT response to sugars and WGA-incubated PC36-SWNT response to sugars. No selective response to WGA conjugate sugars is observed, unlike with ProLoop-SWNT.

### SI. References


(1) Bonis-O'Donnell, J. T. D.; Page, R. H.; Beyene, A. G.; Tindall, E. G.; McFarlane, I. R.; Landry, M. P. Dual Near-Infrared Two-Photon Microscopy for Deep-Tissue Dopamine Nanosensor Imaging. *Adv. Funct. Mater.* **2017**, *27* (39), 1–10.

(2) Nam, K. T.; Shelby, S. A.; Choi, P. H.; Marciel, A. B.; Chen, R.; Tan, L.; Chu, T. K.; Mesch, R. a; Lee, B.-C.; Connolly, M. D.; et al. Free-Floating Ultrathin Two-Dimensional Crystals from Sequence-Specific Peptoid Polymers. *Nat. Mater.* **2010**, *9* (5), 454–460.

(3) Zhao, R.; So, M.; Maat, H.; Ray, N. J.; Arisaka, F.; Goto, Y.; Carver, J. A.; Hall, D. Measurement of Amyloid Formation by Turbidity Assay—seeing through the Cloud. *Biophys. Rev.* **2016**, *8* (4), 445–471.

(4) Armbruster, D. A.; Pry, T. Limit of Blank, Limit of Detection and Limit of Quantitation. *Clin. Biochem. Rev.* **2008**, *29 Suppl 1* (August), S49-52.
